# Evaluating discordant somatic calls across mutation discovery approaches to minimize false negative drug-resistant findings

**DOI:** 10.1101/2023.10.26.562640

**Authors:** Hsin-Fu Lin, Pei-Miao Chien, Chinyi Cheng, Tzu-Hang Yuan, Yu-Bin Wang, Pei-Lung Chen, Chien-Yu Chen, Jia-Hsin Huang, Jacob Shujui Hsu

**Author notes:** Corresponding author, No.1 Jen Ai road section 1, Taipei, 100233, Taiwan, R.O.C. Hsin-Fu Lin and Pei-Miao Chien, contributed equally to this study as co-first authors.

## Abstract

Evaluating robustness of somatic mutation detections is essential when utilizing whole exome sequencing (WES) for treatment decision-making. A comprehensive evaluation was conducted using tumor WES from the FDA-led Sequencing Quality Control Phase 2 (SEQC2) project, in which multiple library kits sequenced identical DNA materials across three labs to benchmark analytical validity. These workflows included various read aligner (BWA, Bowtie2, DRAGEN-Aligner, DRAGMAP, and HISAT2) and mutation caller (Mutect2, TNscope, DRAGEN-Caller, and DeepVariant) combinations. The results revealed that DRAGEN exhibited superior performance, achieving mean F1-scores of 0.966 and 0.791 for SNV and INDEL detection, respectively. Among open-source software, BWA Mutect2 and HISAT2 Mutect2 combinations showed the highest mean F1-scores for SNV (0.949) and INDEL (0.722), respectively. The analyses indicated that high-quality data can be analyzed as having worse results, and vice versa. Evaluations of COSMIC reported mutations unveiled discrepancies across enrichment kits. IDT enrichment kits showed a higher false negative rate, while Agilent WES kits tended to miss mutations in *CBL* and *IDH1*, and Roche library kits tended to miss the mutations in *PIK3CB*. For drug-related biomarkers, Sentieon TNscope tended to underestimate tumor mutation burden and overlook crucial drug-resistance mutations such as *FLT3* (c.G1879A: p.A627T) for cytarabine resistance in leukemia and *MAP2K1* (c.G199A:p.D67N) for *BRAF* inhibitors in melanoma. The findings highlight the importance of robust bioinformatic analysis in identifying tumor mutations and guiding clinical decision-making.

**Highlights:**

- Mutation callers had a significantly higher effect on overall sensitivity than aligners.
- Benchmarking analyses demonstrated that high-quality sequencing reads can be analyzed as having worse results, and vice versa.
- DRAGEN exhibited the best performance among other aligner-caller combinations.
- The combination of BWA with Mutect2 and HISAT2 with Mutect2 yielded the highest mean F1 scores for detecting SNVs and INDELs by open-source software, respectively.
- Sentieon TNscope tended to underestimate the tumor mutation burden and missed several drug-resistant mutations.

## 1. Introduction

Somatic mutations are pivotal for understanding tumor cell growth and providing molecular information for clinical oncology. Several cancer treatments, such as targeted therapies, rely heavily on identifying specific somatic mutations, such as those in *BRAF, VEGF, EGFR*, and *BRCA*, across different cancer types. Moreover, numerous somatic mutations such as those in *EGFR, FLT3*, and *MAP2K1* are associated with drug resistance, indicating the importance of mutation detection. Conflicting results in mutation analyses could result in different downstream clinical decisions. Tumor-specific mutations can be detected by sequencing formalin-fixed paraffin-embedded (FFPE) tissues from tumor lesions. Somatic mutations can also be detected in peripheral blood DNA, as tumor DNA is released into the circulating system as circulating tumor DNA (ctDNA). An increasing number of studies have shown that surveillance of somatic mutations in ctDNA can indicate in situ tumor status. The identification of clinically informative mutations in ctDNA is dependent on a robust analytical workflow. The variant allele frequency (VAF) of a germline genetic variant is around 50%, whereas that of a somatic mutation (which is correlated with the proportion of tumor cells) in ctDNA can be as low as 1%, making it difficult to detect the mutation signal from background noise. Compared to germline genetic variants, callers must be highly sensitive to somatic variants, which often have relatively low allele frequencies. High-throughput sequencing holds promise for detecting low VAF mutations by increasing the sequencing depth. Instead of detecting single nucleotide variants (SNVs), in difficult-to-map or repetitive genomic regions, read mapping can be ambiguous, posing challenges in determining the accurate position of small insertions/deletions (INDELs)[1, 2, 3]. Despite the availability of several successful aligners and callers, their reproducibility remains challenging. As different callers can report the same INDEL in various ways, variant format normalization is recommended to compare different callsets. Therefore, systematic validation of analytical workflows is necessary.

The lack of standardized biological specimens often leads to inconclusive information in the field. Unlike germline variations based on several individual samples in the genome in a bottle (GIAB) consortium, benchmarking materials for somatic mutation detection often rely on synthetic or simulated datasets. Recently, the Food and Drug Administration-led sequencing and quality control phase 2 (SEQC2) consortium released a set of benchmarking tumor cell WES data (Supplementary Table S1) for somatic mutation detection, especially for the tumor-only mode[4, 5], a focus of this study. Benchmarking materials including raw sequencing reads, ground truth mutation calls and confidence genomic regions, provide a gold standard for clinical investigations and improve data reproducibility. Consequently, the evaluation of different somatic variant detection tools has become feasible. Benchmarking materials including raw sequencing reads, ground truth mutation calls and confidence genomic regions, provide a gold standard for clinical investigations and improve data reproducibility. Consequently, the evaluation of different somatic variant detection tools has become feasible.

Recent studies[6, 7] have demonstrated that the use of whole exome sequencing (WES) or whole genome sequencing (WGS) data can lead to precise cancer treatment. However, previous studies have shown that aligners and variant callers can notably influence the results of variant detection[8, 9]. Additionally, several studies have indicated that different tool combinations often generate discordant variant detection results[10, 11, 12, 13, 14, 15]. In this study, various combinations of aligners and callers were compared. Several reputable aligners and callers were selected and their performance was evaluated. The following aligners were selected: BWA version 202112(Sentieon built in version)[16], Bowtie2 version 2.4.2[17], Illumina DRAGMAP version 1.3.0 (https://github.com/Illumina/DRAGMAP, accessed 26 July 2024.), the built-in aligner of the Illumina commercialized DRAGEN version 07.021.645.4.0.3 Bio-IT Platform (abbreviated as DRAGEN-A) https://www.illumina.com/products/by-type/informatics-products/dragen-secondary-analysis.html, accessed 26 July 2024. The software was provided by Illumina (5200 Illumina Way San Diego, CA 92122 USA.), and the graph-based aligner HISAT2 version 2.2.0[18]. The callers for the cancer variants included GATK Mutect2 verion 4.2.3.0 [19], Sentieon TNscope verson 202112 [20], Illumina DRAGEN version 07.021.645.4.0.3 Bio-IT Platform built-in caller (DRAGE-NC), and Google DeepVariant version 1.4.0[21]. Notably, Sentieon TNscope was designed based on the Mutect2 algorithm with improved speed. Detailed information on all aligners and callers is provided in Supplementary Figure S1/Table S2. The SEQC2 benchmarking dataset was reanalyzed, and 18 combinations of aligners and callers were compared for their performance in the tumor-only mode for detecting cancer single nucleotide variants (SNVs) and small INDELs. The results provide critical information and show that no single combination produces superior results for detecting all variants in samples. In other words, none of these tools and combinations can have a detection rate of 100%. A merged or ensemble-based approach, which combines the results from multiple combinations, is most likely to improve the performance of variant detection in tumor samples.

## 2. Materials and Methods

### 2.1. Benchmark

The benchmarking materials comprised nine replicates of paired-end whole exome sequencing (WES) reads derived from the same biological sample, utilizing a mixture of cancer cell lines developed by the SEQC2 consortium[4]. The same biological replicates were used across three different commercial WES library preparation kits in three laboratories. The ground truth datasets includes verified mutation calls for 41,395 SNVs, 537 INDELs, and 266 multinucleotide variants (MNVs) in the confidence genomic regions. 52.17% of the truth positive mutations had a VAF lower than 0.2, providing a gold standard for clinical oncology investigations and enhancing reproducibility. The benchmarking dataset and confidence regions were downloaded SRA (Accession numbers: SRR13076390 to SRR13076398), and they are referred to as S1–S9 in this study. The nine sets of paired-end WES reads from three different exome enrichment kits, including 1) the Agilent kit (Agilent Technologies, Santa Clara, CA), 2) KAPA Hyper Prep kit (Kapa Biosystems) plus Roche NimbleGen SeqCap EZ hybridization and wash kit (Roche Sequencing Solutions, Indianapolis, IN) with Next Ultra II DNA Library Prep kit for Illumina (New England Biolabs, Ipswich, MA), and 3) IDT xGen hybridization and wash kit (Integrated DNA technologies, Coralville, Iowa). Each kit had three technical replicates conducted by three different laboratories. Details of the nine samples are presented in Supplementary Table S1.

### 2.2. Aligner and variant caller combination implementations

The entire analytical workflow of this study is illustrated in Supplementary Fig. S1, and the tool versions are listed in Supplementary Table S2. Five aligners and four callers were adopted, forming up to 20 different combinations. ALIGNER CALLER was used to represent the different combinations for a callset. For example, DRAGEN-A DRAGEN-C represents the Illumina-commercialized DRAGEN Bio-IT Platform for aligner and caller. The first phase involved processing raw reads to produce analysis-ready BAM files. This included read alignment against the GATK human g1k v37 decoy.fasta reference genome followed by read deduplication. Forty-five (five aligners × nine samples) text-based Sequence Alignment Map files were obtained and converted into the analysis-ready binary version format files (BAM). The second phase was variant discovery from analysis-ready BAM files and left-normalization using bcftools in the VCF format. However, HISAT2 DRAGEN-C and HISAT2 DeepVariant encountered data formatting problems, resulting in their exclusion from the analysis. Thus, 18 different combinations were evaluated.

### 2.3. Performance evaluation

GATK SelectVariants was used to filter out variants that were not in the confidence regions and compared them with the benchmark using Haplotype Comparison Tools (https://github.com/Illumina/hap.py, accessed 26 July 2024.) and programs developed in-house to evaluate the performance of the combinations. The performance of each tool combination was evaluated by comparing the VCF file against the SEQC2 truth set benchmark VCF file and calculating the F1 score, recall, and precision. The F1 score was calculated as follows:

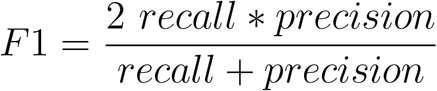

### 2.4. Computing environment and resources

All analyses were performed using the high-performance computing (HPC) cluster computing environment Taiwania 3 at the National Center for Highperformance Computing, Taiwan. For DeepVariant, the Docker image was obtained from https://github.com/google/deepvariant, accessed 26 July 2024. and installed on a Google Kubernetes Engine (GKE)-based HPC cluster computing environment in collaboration with Taiwan AI Labs.

### 2.5. Analysis of COSMIC gene census and cancer-related biomarkers

Exon regions were defined by SEQC2, which contained 22,519,561 base regions. Curated cancer gene censuses (N = 738) and drug-associated genes (N = 15) were obtained from the Catalogue of Somatic Mutations in Cancer (COSMICv3.3)[22]. The SEQC2 truth set contained 512 genes (512/738; 69.3%) of cancer gene census. In addition, the TMB and mutational signatures were analyzed. The TMB was calculated as follows:

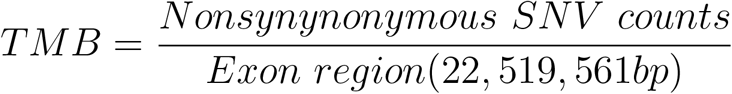

Mutational signature analyses were performed with SigProfilerExtractor v1.1.4 (https://github.com/AlexandrovLab/SigProfilerExtractor, accessed 26 July 2024.) using the latest COSMIC v3.3 signature database.

### 2.6. Data availability

The data was downloaded from SEQC2, released in 2021 (SRA download link is https://www.ncbi.nlm.nih.gov/sra, accessed 26 July 2024. Besides, SRA number is SRP292966 and accession numbers is SRR13076390 to SRR13076398).

### 2.7. Code availability

The underlying codes for this study are available in Github https://github.com/Jacob-s-Lab/Somatic_pipeline_JMD, accessed 31 July 2024.

## 3. Results

### 3.1. Comparison of different WES library preparation kits

The average mapping depth of nine samples were evaluated to analyze deduplicated alignments using Picard (version 2.27.4). Regardless of the aligners used, distinct patterns were observed in the library preparation methods. Samples treated with the IDT WES enrichment kit exhibited lower mapping depths and higher duplication rates (Supplementary Table S3), whereas those treated with the Agilent kit showed higher mapping depths.

Among the nine samples analyzed, S3 (SRR13076392) consistently exhibited the highest mapping depth and lowest duplication rate, whereas S4 (SRR13076393) consistently showed the opposite quality. Among the same Agilent library preparation kits, even though the data from laboratory 3 (S3 dataset) generated approximately 16.5% more reads than the data from laboratory 1 (S1 dataset), they achieved a lower duplication rate with at least 50% more depth. These observations highlight the importance of considering both library preparation methods and sample quality when interpreting results. In addition, when comparing the results obtained from the five aligners, they yielded similar mapping depths with minimal differences for each sample (Supplementary Fig. S2). The depth analysis results suggested that the Agilent samples had a higher mapping average depth and the IDT samples had the lowest mapping average depth.

Depth comparison results indicated that the choice of aligner has a limited effect on the observed mapping depth. Overall, these findings emphasize the significance of library preparation methods and sample quality in influencing mapping depth and duplication rates, and suggest that the choice of the aligner may have a moderate effect.

### 3.2. Comparison of the overall SNV and INDEL calling abilities

To obtain an overview of the performance of all variant calling and alignment combinations, the F1-score of 18 different combinations was emohasized. The results were separated into SNVs (Figure 2a, Supplementary Figures S3 and S4, Supplementary Table S4) and INDELs (Figure 2b, Supplementary Figures S3 and S4, Supplementary Table S5).

**Figure 1:**
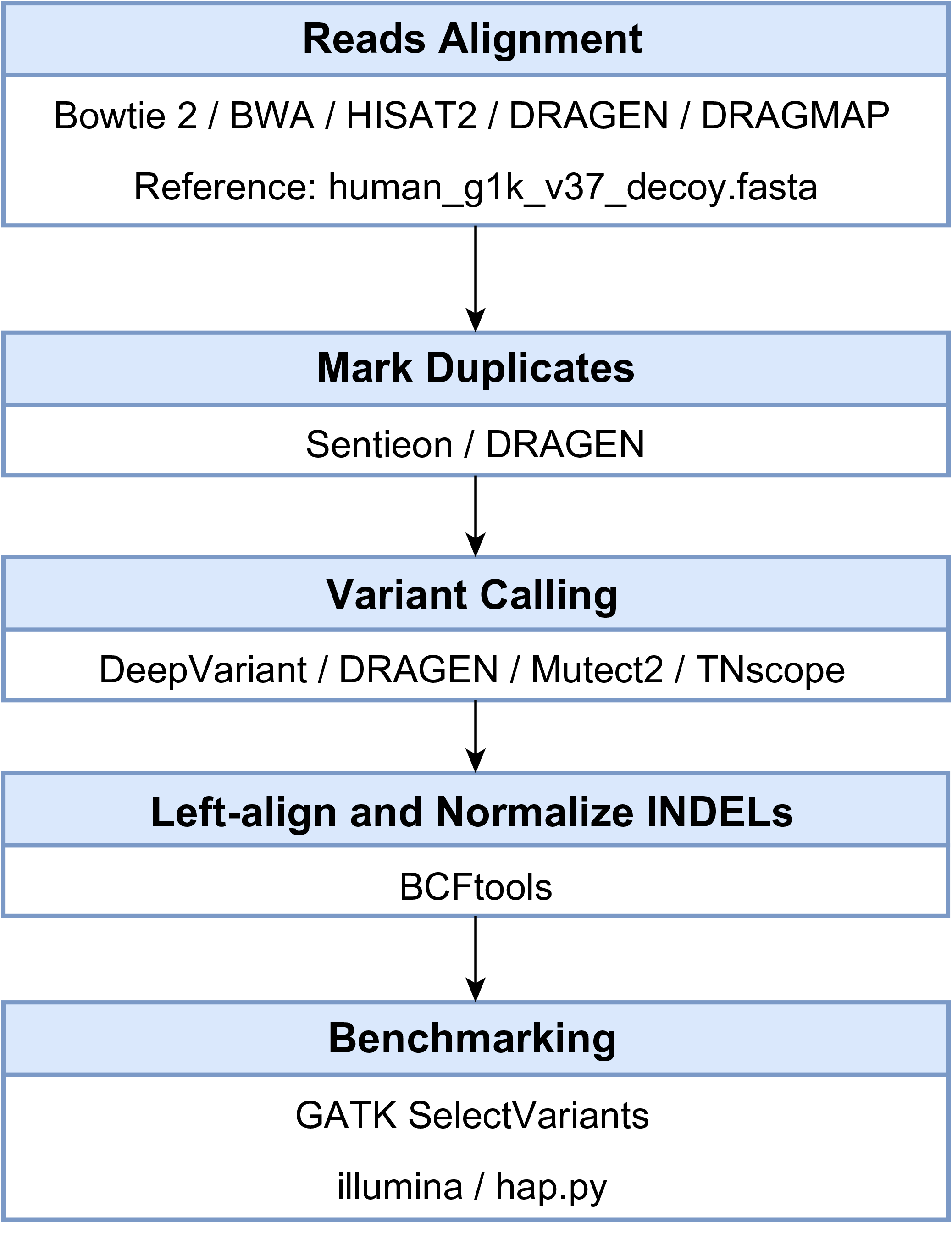
Workflow in this study and tools used in every step.

**Figure 2:**
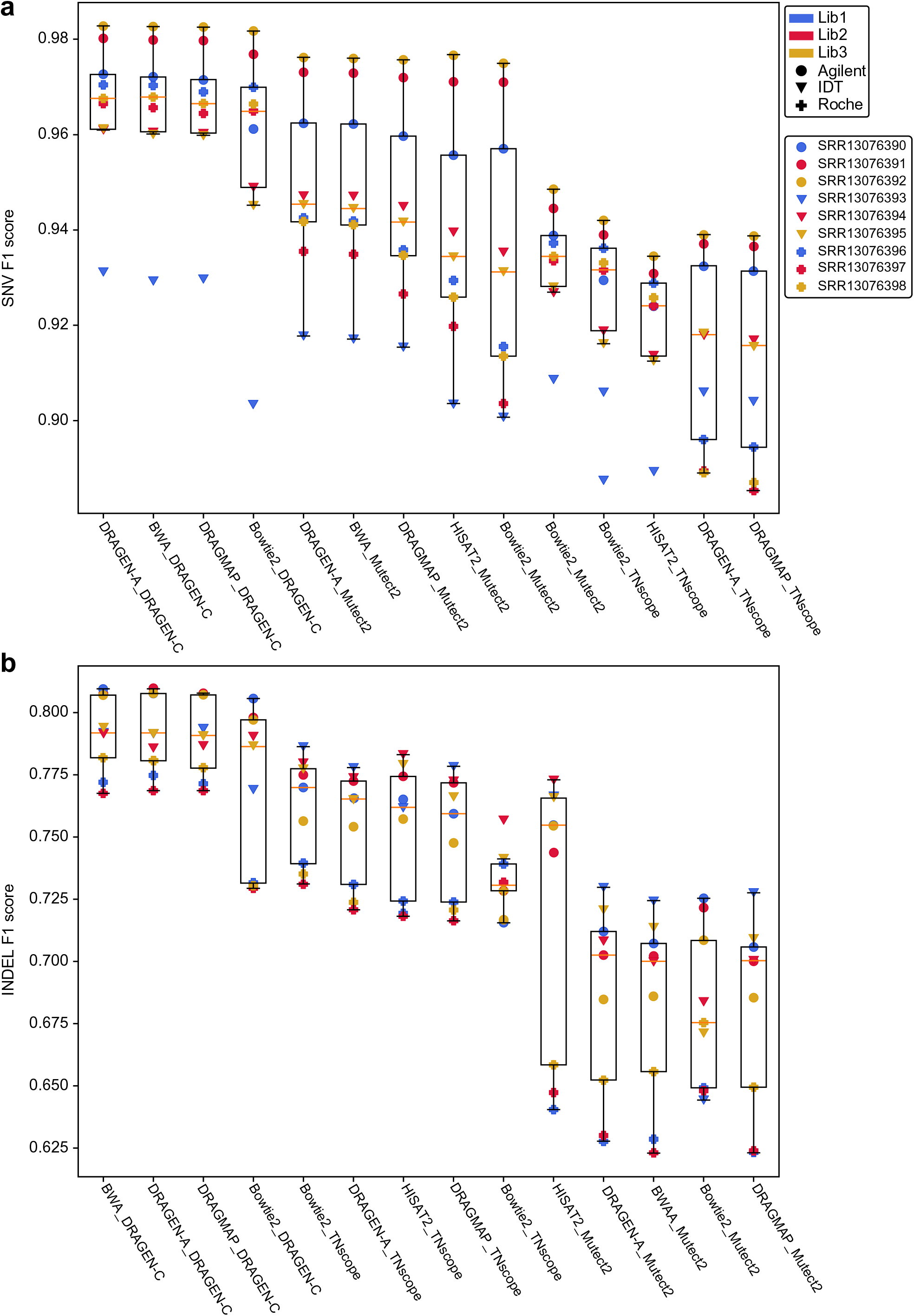
Comparison of the F1 scores of somatic mutation calls among workflows with SEQC2 WES datasets as the truth set. Comparison of the overall F1 scores for the nine WES samples among three different labs and three library preparation kits (Agilent/IDT/Roche) under different workflows for (a) SNVs and (b) INDELs.

For SNV detection, the built-in DRAGEN, a combination of DRAGEN-A and DRAGEN-C, showed the best performance, achieving a mean F1-score of 0.966, mean recall rate of 0.954, and mean precision of 0.979 (Table 1, Supplementary Table S4). Among all the open-source combinations, BWA Mutect2 had the best performance in terms of SNV, with a mean recall rate of 0.953, mean precision of 0.944, and mean F1-score of 0.949. It was observed that sample 3 (SRR13076392) from laboratory 3 showed the highest F1 scores, followed by sample 2 from laboratory 2 and sample 1 from laboratory 1, indicating variant call inconsistencies introduced by the laboratories.

**Table 1:** Recall, precision, and F1 score of the SNV and INDEL results under different combinations. The table only includes the best combination for commercial tools, opensource combination, and worst. The numbers indicate the recall/precision/F1 scores.

| TYPE | SNV |  | INDEL |  |
| --- | --- | --- | --- | --- |
| — | Commercial | Open-source | Commercial | Open-source |
| Combination | DRAGEN-A | BWA | DRAGEN-A | HISAT2 |
| Sample No. | _DRAGEN-C | _Mutect2 | _DRAGEN-C | _Mutect2 |
| SRR13076390 | 0.964/0.982/0.973 | 0.956/0.968/0.962 | 0.970/0.692/0.808 | 0.879/0.661/0.755 |
| SRR13076391 | 0.982/0.978/0.980 | 0.976/0.970/0.973 | 0.983/0.688/0.810 | 0.910/0.629/0.744 |
| SRR13076392 | 0.990/0.976/0.983 | 0.983/0.969/0.976 | 0.991/0.682/0.808 | 0.922/0.639/0.754 |
| SRR13076393 | 0.883/0.985/0.931 | 0.889/0.947/0.917 | 0.878/0.721/0.792 | 0.807/0.730/0.767 |
| SRR13076394 | 0.940/0.983/0.961 | 0.940/0.954/0.947 | 0.941/0.675/0.786 | 0.881/0.688/0.773 |
| SRR13076395 | 0.942/0.982/0.961 | 0.942/0.947/0.944 | 0.931/0.689/0.792 | 0.855/0.693/0.766 |
| SRR13076396 | 0.967/0.974/0.970 | 0.967/0.918/0.942 | 0.970/0.645/0.775 | 0.892/0.499/0.640 |
| SRR13076397 | 0.958/0.975/0.966 | 0.964/0.907/0.935 | 0.965/0.639/0.769 | 0.891/0.508/0.647 |
| SRR13076398 | 0.960/0.975/0.968 | 0.964/0.920/0.941 | 0.961/0.657/0.781 | 0.885/0.524/0.658 |
| Average | **0.954/0.979/0.966** | 0.953/0.944/0.949 | 0.954/0.676/0.791 | 0.880/0.619/0.723 |

**Table 2:** Mutational signature results for S3/S4 samples in DRAGEN, BWA_Mutect2, and Bowtie2_DeepVariant. Details for each sample are provided in Supplementary Table S10.

|  | Aligner | Caller | SBS (N = 96) | DBS (N = 11) | ID (N = 18) |
| --- | --- | --- | --- | --- | --- |
| Know true set | — | — | SBS1 (17.58%) | DBS10 (64.42%) | ID4 (30.46%) |
|  |  |  | SBS5 (51.32%) | DBS11 (35.58%) | ID15 (32.7%) |
|  |  |  | SBS6 (18.16%) |  | ID16 (36.84%) |
|  |  |  | SBS54 (12.94%) |  |  |
| S3 | DRAGEN-A | DRAGEN-C | SBS1 (17.38%) | DBS10 (64.04%) | ID7 (46.04%) |
|  |  |  | SBS5 (51.42%) | DBS11 (35.96%) | ID10 (27.64%) |
|  |  |  | SBS6 (18.36%) |  | ID16 (26.32%) |
|  |  |  | SBS54 (12.84%) |  |  |
|  | BWA | Mutect2 | SBS1 (17.20%) | DBS10 (60.94%) | ID7 (57.82%) |
|  |  |  | SBS5 (51.42%) | DBS11 (39.06%) | ID10 (42.18%) |
|  |  |  | SBS6 (18.42%) |  |  |
|  |  |  | SBS54 (12.96%) |  |  |
|  | Bowtie2 | DeepVariant | SBS1 (19.28%) | DBS10 (100%) | ID4 (29.5%) |
|  |  |  | SBS5 (58.32%) |  | ID15 (29.32%) |
|  |  |  | SBS54 (22.40%) |  | ID16 (45.18%) |
| S4 | DRAGEN-A | DRAGEN-C | SBS1 (17.56%) | DBS10 (63.42%) | ID7 (41.96%) |
|  |  |  | SBS5 (51.40%) | DBS11 (36.58%) | ID10 (31.02%) |
|  |  |  | SBS6 (17.06%) |  | ID16 (27.02%) |
|  |  |  | SBS54 (13.98%) |  |  |
|  | BWA | Mutect2 | SBS1 (17.38%) | DBS10 (60.86%) | ID7 (48.18%) |
|  |  |  | SBS5 (52.48%) | DBS11 (39.14%) | ID10 (51.82%) |
|  |  |  | SBS6 (16.66%) |  |  |
|  |  |  | SBS54 (13.48%) |  |  |
|  | Bowtie2 | DeepVariant | SBS1 (19.50%) | DBS10 (100%) | ID7 (33.0%) |
|  |  |  | SBS5 (58.58%) |  | ID10 (29.12%) |
|  |  |  | SBS54 (21.92%) |  | ID16 (37.88%) |

Among all the combinations, sample 3 (SRR13076392) was always the best, whereas sample 4 (SRR13076393) was always the worst. However, for sample 3 (SRR13076392), some analysis combinations led to worse results compared with those for sample 4 (SRR13076393) with proper analyses. This finding highlights the importance of tool selection and data analysis. High-quality data can be analyzed as having worse results, whereas low-quality data can still achieve relatively better results. Overall, the Agilent WES library preparation kit had higher F1 scores than other kits. However, although the library preparation kits were identical, the F1 scores varied, indicating the importance of benchmarking for variant calling analyses. Sample 3 (SRR13076392) was used for all downstream analyses to minimize the effects of library preparation and sequencing. Notably, all of the combinations with DeepVariant had relatively poor performance with F1, recall, and precision of 0.4, 0.3, and 0.6, respectively, for SNVs, and 0.3, 0.2, and 0.5, respectively, for INDELs (Table 1, Supplementary Table S4), indicating that DeepVariant is unsuitable for tumor-only somatic mutation detection. Therefore, DeepVariant was excluded from further analysis. In general, each sample showed a similar pattern in the 14 analytical workflows, indicating the importance of library preparation.

For small INDEL identification, BWA DRAGEN-C had the best performance, slightly better than the built-in DRAGEN, with a mean F1-score of 0.792, recall rate of 0.954, and precision of 0.678. Among all open-source combinations, the HISAT2 Mutect2 combinations had the highest mean F1 score (0.72) with a mean recall of 0.88 and mean precision of 0.62. In contrast, DRAMAP Mutect2 had the highest mean recall rate (0.95) with a mean precision of 0.53 and mean F1 score of 0.68 (Supplementary Table S5), indicating that HISAT2 Mutect2 is a more balanced combination by sacrificing detection sensitivity. Unlike the SNV detection patterns, it was observed that commercial variant callers tended to perform better, with DRAGEN-C being the best, followed by TNscope. Sample 3 (SRR13076392) was never the best dataset for INDEL detection. The Agilent WES library preparation kits performed better for DRAGEN-C and the IDT WES kits performed better for the other callers. It was observed that varied laboratory performances even with identical library preparation kits.

It was found that the selection of analysis tool combinations influenced the variant calling results, as suitable analysis tools may compensate for low mapping depth sample accuracy. For example, IDT WES kit samples 5 and 6 had a relatively low depth of coverage; however, the overall SNV F1 scores in DRAGEN were higher than those of many other samples. It outperformed all sample results in the TNscope group. In addition, although sample 4 (SRR13076393) had the lowest depth of coverage, its INDEL calling performance exceeded that of the other high-depth coverage samples. For Mutect2, calling the INDEL group, sample 4 (SRR13076393) was the best among different samples.

Variant calls among the aligners and callers were also dissected. The variant call was calculated for the same aligner with different callers and vice versa. The analysis revealed that variant callers played a more critical role than aligners in the overall process. Specifically, approximately 94% of the variants were observed at the intersection of the five aligners when the same caller was used. However, when using the same aligner, only approximately 40% of the variants overlapped among the four callers (Supplementary Figures. S5–S8). Of all the callers analyzed, Mutect2 had the highest intersection with DRAGEN-C, indicating that its results were similar to those produced by DRAGEN-C.

### 3.3. Combining different tool combinations can improve the overall detection capability

As no single tool can achieve perfect somatic mutation detection, to achieve an F1 score higher than that with existing tools, the possibility of merging the results from various combinations was systematically evaluated. The approach involves only open-source tools, with the exception of DeepVariant. It was found that combining multiple callsets can improve the overall performance. For SNV, merging Mutect2 with different aligners, such as BWA HISAT2 and DRAGMAP, led to slight improvements in the highest F1 score of 0.9779 (Figure 3a, Supplementary Table S6), which is very close to that of DRAGEN at 0.9827. For INDEL, the highest F1 score for a single combination was for DRAGEN (DRAGEN-A DRAGEN-C) at 0.8078 and that for the open-source combination was for HISAT Mutect2 at 0.6506. However, several merging approaches outperformed DRAGEN (Figure 3b, Supplementary Table S7). For example, merging results from Bowtie2 Mutect2 and BWA TNscope can increase the F1 score to 0.8126. Our findings underscore the value of combinations, demonstrating enhanced overall performance compared to the best opensource combination BWA Mutect2 for SNVs and HISAT2 Mutect2 for IN-DELs. However, DRAGEN as a single tool remained the best.

**Figure 3:**
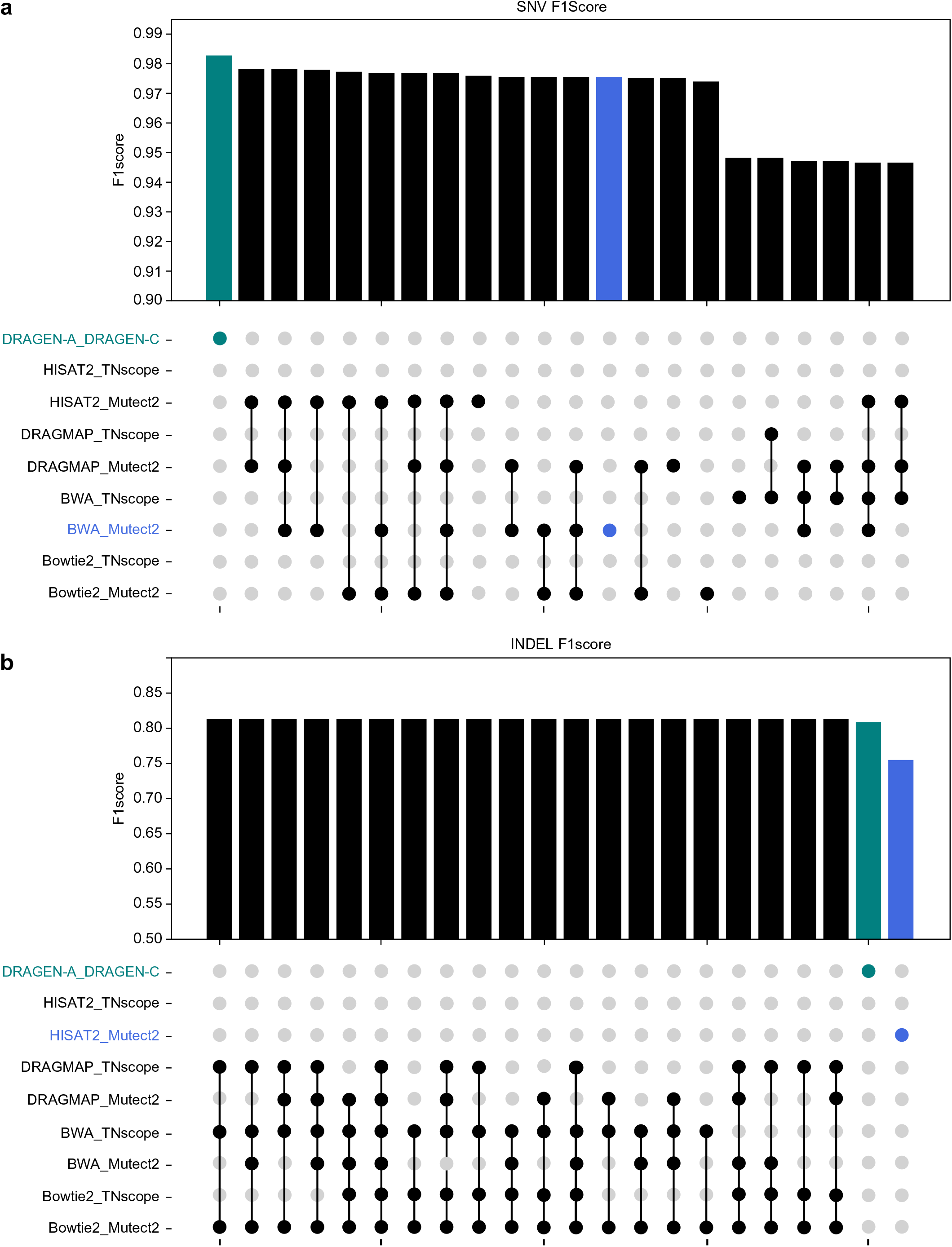
Top 20 F1 scores of different workflow coalitions: (a) SNVs and (b) INDELs. The green bar indicates the performance of the built-in DRAGEN (DRAGEN_A_GRAGEN_C) and the blue bar represents the best open-source combination, BWA_Mutect2 for SNV and HISAT2_Mutect2 for INDEL.

### 3.4. Limitation of detecting the lowest VAF

To investigate the performance of different variant calling tools in detecting somatic variants within a spectrum of VAF, the VAFs of the truth set were divided into a range of 100 bins. In the truth set, most of the variants were distributed in the VAF *<* 0.2 region (Figure 4a). Using the best-quality sample 3 (SRR13076392), the best callset of all variant calling tools was systematically dissected. It was found that DRAGEN-C, Mutect2, and TNscope shared the same VAF detection limit of 0.0063, whereas the lowest VAF in the truth set was 0.0058. Subsequently, all true positives (TPs), false negatives (FNs), and false positives (FPs) across the VAF spectrum were compared (Figure 4b).

**Figure 4:**
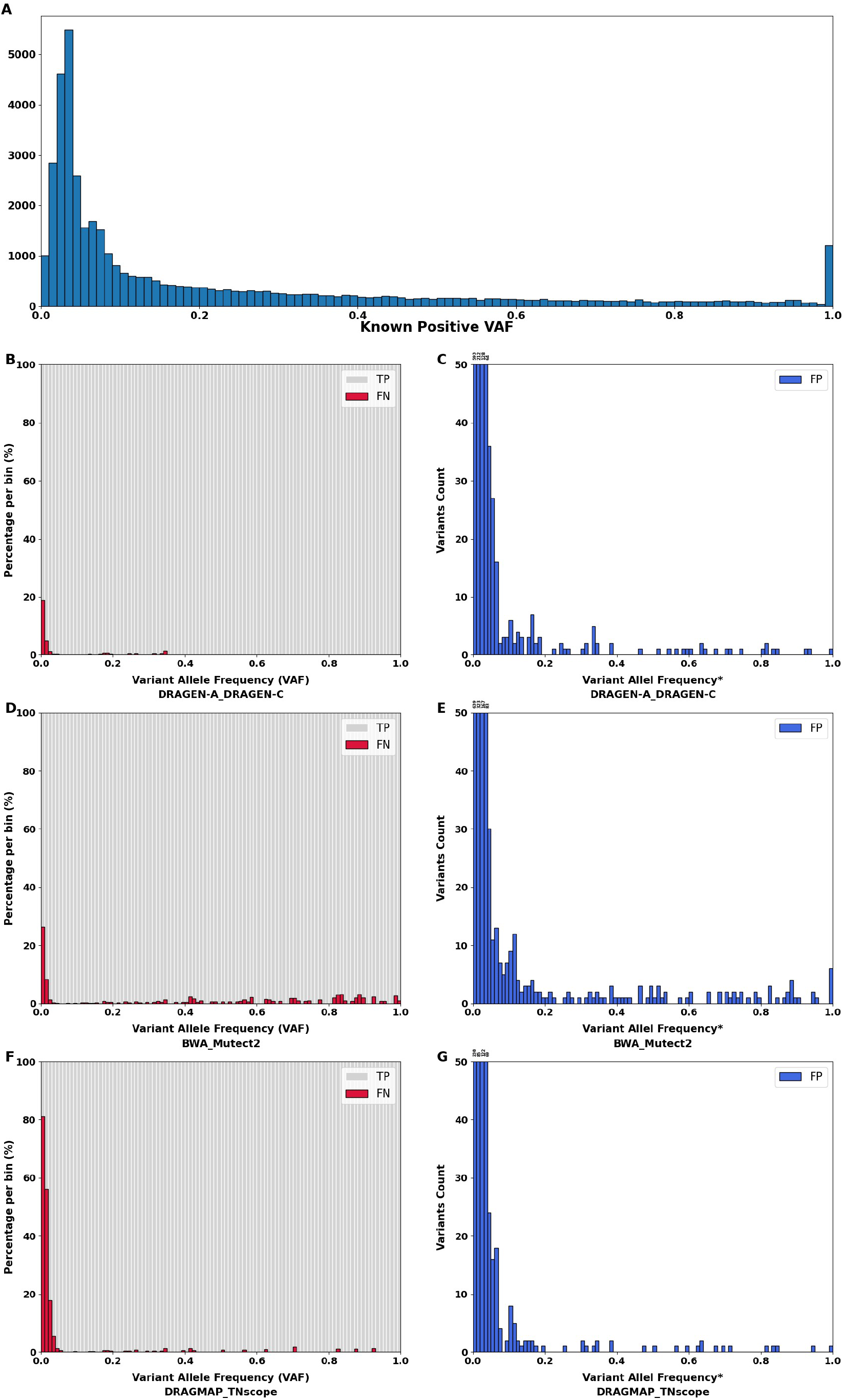
Variant allele frequency (VAF) spectra for truth set and the calling results from selected combinations. The results were based on the S3 (SRR13076392) dataset for the best workflow (DRAGEN-A_DRAGEN-C) and two best open-source workflows (BWA_Mutect2 for SNV and DRAMAP_TN-scope for INDEL). (A) VAF distribution in the SEQC2 benchmark truth set. (B, D, and F) The red bar represents the percentage of false negative (FN) calls and the grey bar is for true positive (TP). (C, E, and G) The blue bar shows the number of false positive (FP) calls across different VAFs. The VAFs are divided into 100 bins from 0 to 1. *VAFs were calculated by each variant calling tool.

DRAGEN-C demonstrated higher sensitivity in detecting most variants, albeit with a small proportion of FNs in extremely low-VAF mutations. Conversely, the BWA Mutect2 combination tended to miss variants across all VAF ranges, suggesting that VAF was not the primary factor for FNs. DRAMAP TNscope displayed a higher rate of FNs, particularly in detecting low-VAF variants, with more than three- to four-fold fewer VAF variants remaining undetected. These findings underscore the distinct capabilities of each variant-calling tool in detecting somatic variants within specific VAF ranges. Regarding FPs, our results indicated that the mutation-detecting tools produced FP calls across the entire range of VAFs, with a considerable proportion of FPs in the low VAF regions. Furthermore, the limitations of merging sample sets and analytical workflows are explored. Combining all the nine samples and all combination results, 42,195 of 42,212 known positive variants (99.95%) were identified, indicating that the current comparisons never detected several TPs.

### 3.5. Effects of tumor mutational burden in different callsets

Tumor mutational burden (TMB) plays a crucial role in oncological genetic testing because of its association with the tumor microenvironment and response to immunotherapy. TMB refers to the total number of somatic mutations in a tumor genome, typically measured as the number of mutations per megabase of sequenced DNA. In this study, the TMB was calculated based on sample and tool combinations and compared to the truth-set TMB, observing tendencies among tools (Supplementary Fig. S9, Table S8). Mutect2 tended to overestimate the TMB in the Roche WES enrichment kit samples. Most of the Agilent & IDT WES enrichment kit samples showed similar TMB values, except for S4 (SRR13076393). The results from Mutect2 imply inconsistent TMB estimations even with identical DNA materials and variant detection analyses, which may lead to different clinical decisions. For DRAGEN-C, most results were closer to the truth set, whereas Bowtie2 DRAGEN-C tended to underestimate the TMB regardless of the WES enrichment kit used. For TNscope, the TMB results varied for different aligners. TNscope tended to considerably underestimate the TMB, except for the Roche enrichment datasets, which may lead to FN TMB results and missed immunotherapy opportunities for patients with cancer. These comparisons emphasize the value of benchmarking analyses, selecting appropriate tools, and library preparation methods in inferring the reliability of TMB results.

### 3.6. Effects of cancer census genes in different callsets

To investigate the performance of different caller variants on cancer-related genes, mutations in a curated cancer census gene list from the COSMIC database (v3.3) were examined. Synonymous SNVs, non-frameshift deletions, and non-frameshift insertions were excluded from the analysis. The most inconsistent results in different combinations were mainly focused on; therefore, only a gene with a FN in at least 11 of the 18 (*>* 60%) combination workflows in any sample was included. The mutation numbers were calculated and compared with the numbers in the truth set as positive discovery results for each sample (Figure 5, Supplementary Table S9). There were 23 genes left; the similarity results are shown in Figure 5. A sample-related issue was considered if all combinations failed to identify true mutations. For example, mutations were located in *CBL* in samples 1, 3, and 6 and in *MSH2* in samples 3, 4, and 6. In DRAGEN-A DRAGEN-C workflows, Figure 6 displays the IGV results for samples 2 and 3 for *CBL* and *MSH2* genes, respectively. Figure 6 (a) showed the TP variant in *CBL* gene in sample 2, but Figure 6 (b) showed the variant was missing in sample 3. Additionally, Figure 6 (c) showed the TP variant in *MSH2* gene in sample 2, but Figure 6 (d) showed the variant was missing in sample 3. Furthermore, Figure 7 (a) showed the TP variant with 1.56% allele frequency in sample 2 (Chr11:119,172,362 C[447] to T[7]), which was larger than the ground truth variant with 1.84% VAF in *CBL* gene. However, Figure 7 (b) shows the variant was missing with 1.06% allele frequency in sample 3 (Chr11:119,172,362 C[563] to T[6]), which was lower than the ground truth variant with 1.84% VAF. Besides, Figure 7 (c) showed the TP variant with 2.36% allele frequency in sample 2 (Chr2:47,637,265 C[538] to T[13]), which was larger than the ground truth variant with 2.35% VAF in *MSH2* gene. However, Figure 7 (d) shows the variant is missing with 0.76% allele frequency in sample 3 (Chr2:47,637,265 C[777] to T[6]), which is lower than the ground truth variant with 2.35% VAF. In a comparison among DRAGEN-C, Mutect2, and TNscope, TNscope tended to exhibit a higher rate of missing detections of these cancer census genes. Library preparation also affected the results, as the patterns from each sample were distinct. Specifically, Agilent WES library kits tended to miss mutations in *CBL* and *IDH1*. The Roche library kits tended to miss the mutation in *PIK3CB*. Samples S4 (SRR13076393), S5 (SRR13076394), and S6 (SRR13076395), which were constructed using IDT enrichment kits, showed a higher rate of FNs than the Roche or Agilent WES library preparation methods. The mutations in *MSH2* were undetected. Overall, our results suggested that different library preparation kits tend to miss mutations on different cancer census genes.

**Figure 5:**
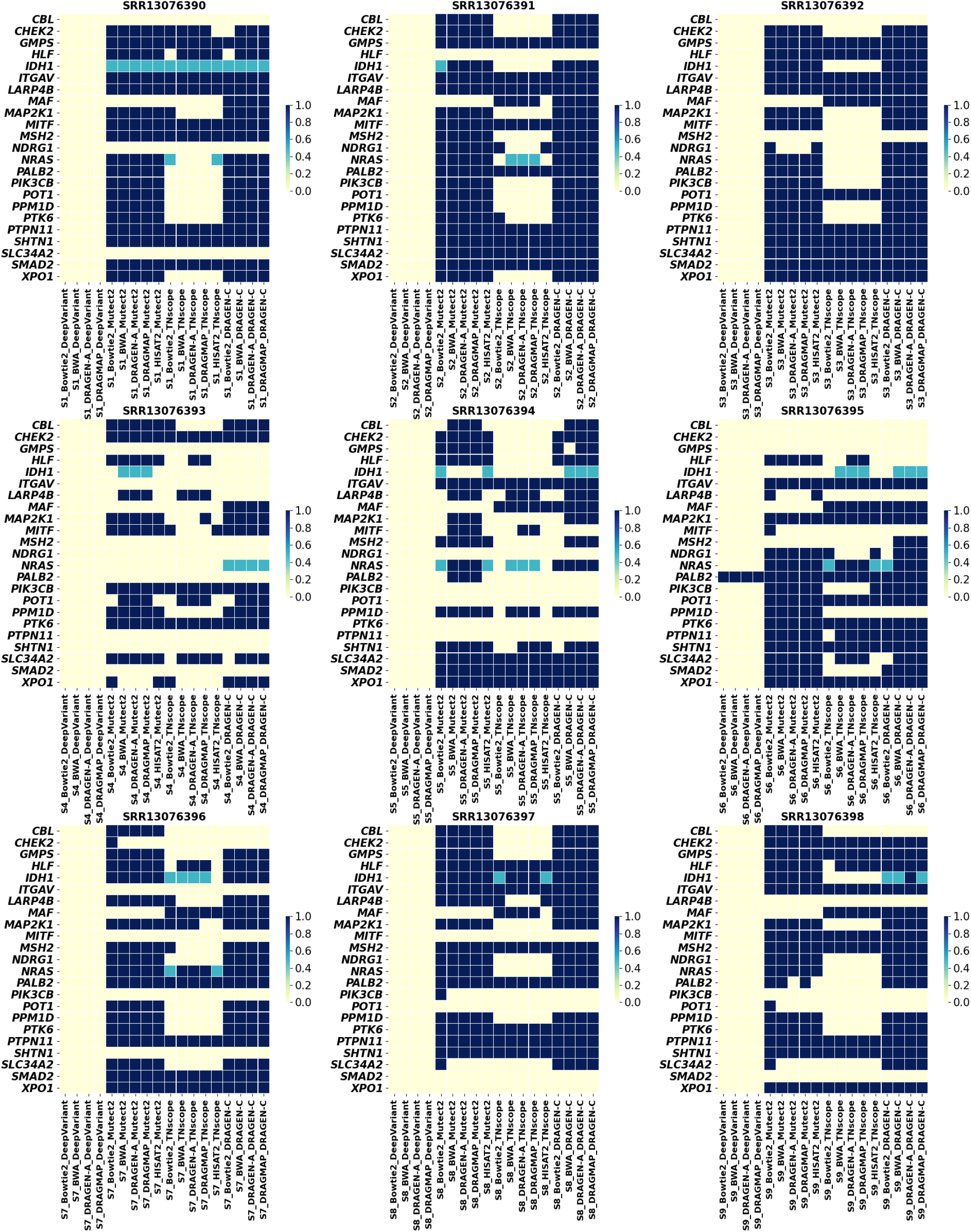
True positive discovery rate of the nine samples under different combinations in cancer census genes.

**Figure 6:**
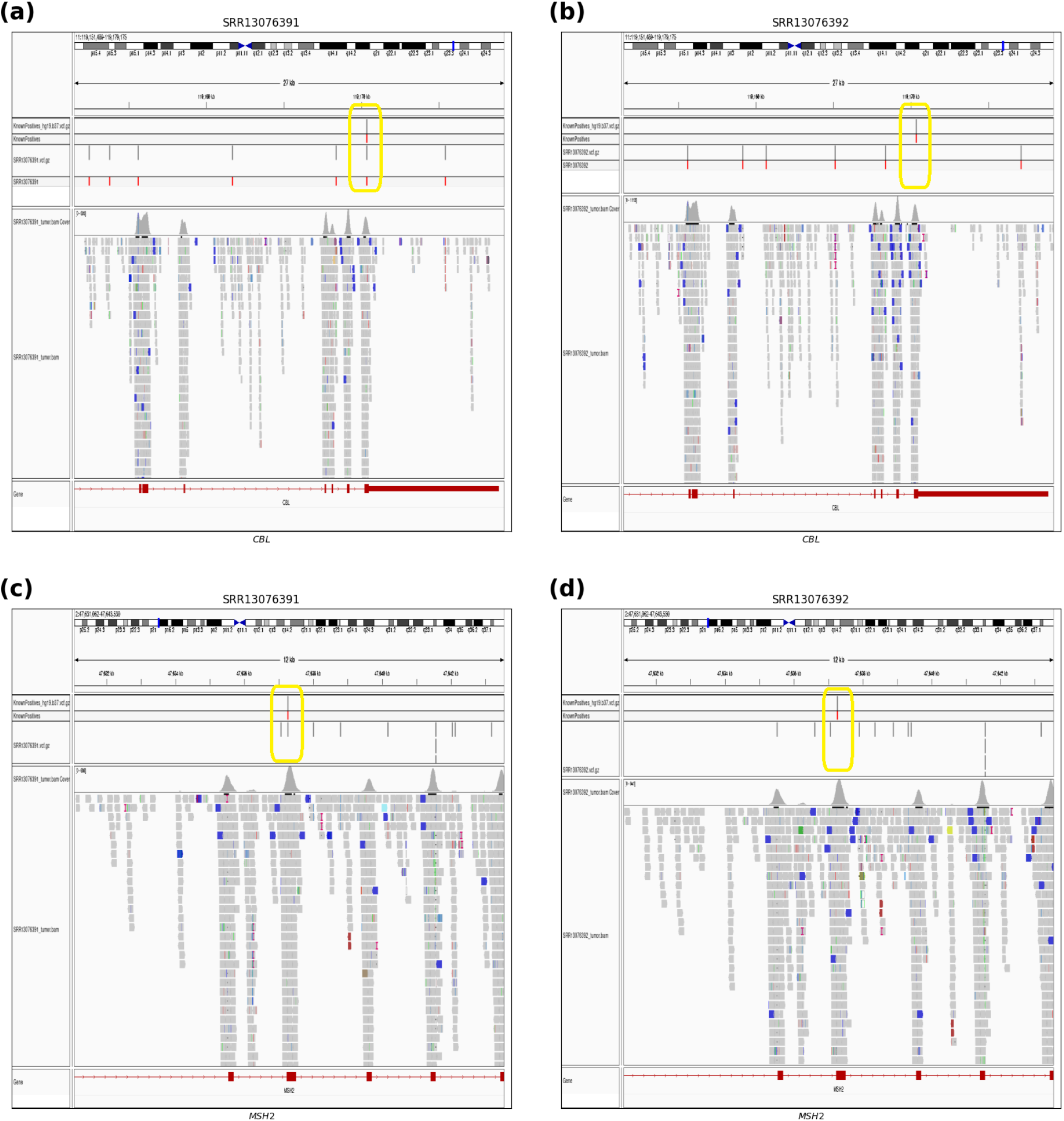
(a) showed the true positive variant in *CBL* gene in sample 2. (b) showed the variant was missing in sample 3. (c) showed the true positive variant in *MSH2* gene in sample 2. (d) shows the variant was missing in sample 3.

**Figure 7:**
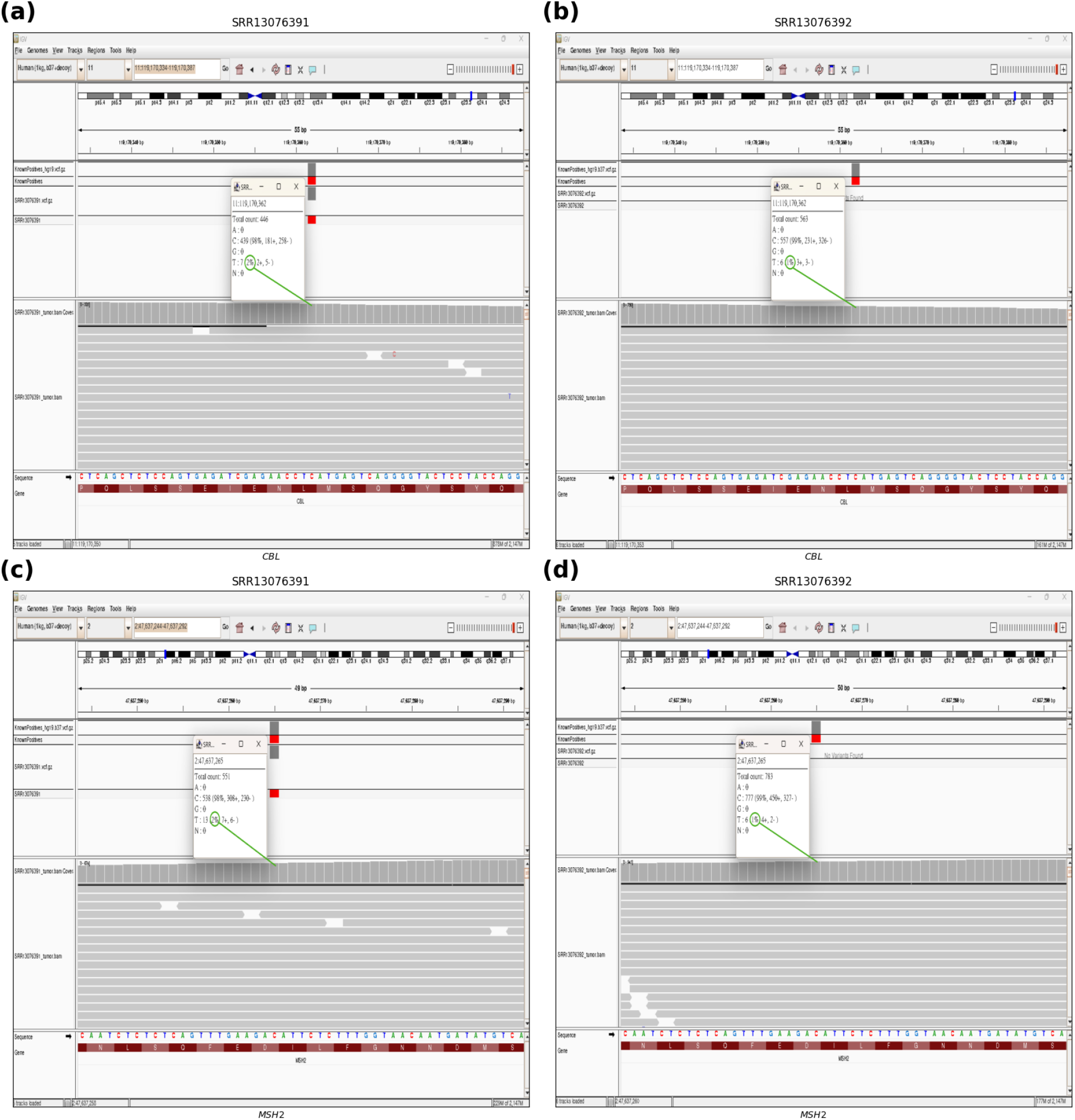
(a) showed the true positive variant with 1.56% allele frequency composed of 447 reference allele C and 7 alternative allele T in sample 2 at site 119,172,362 on chromosome 11 in *CBL* gene. (b) shows the variant was missing with 1.06% allele frequency composed of 563 reference allele C and 6 alternative allele T in sample 3. Furthermore, (c) showed the true positive variant with 2.35% allele frequency composed of 538 reference allele C and 13 alternative allele T in sample 2 at site 47,637,265 on chromosome 2 in *MSH2* gene. However, (d) showed the variant was missing with 0.76% allele frequency composed of 777 reference allele C and 6 alternative allele T in sample 3.

### 3.7. Effects of drug-associated mutations in different callsets

The mutation-detection ability of 15 drug-associated genes listed in the COSMIC database was also checked. *ABL1* and *BTK* were not included in the truth set, and the mutations in *JAK1, JAK2*, and *SMO* were synonymous SNVs. Therefore, these five genes were excluded from the comparison. The focus was on genes that have been documented to be associated with drug resistance or to play a role in treatment selection. The mutation detection capability in these genes may influence decision-making for personalized treatment. Our analyses revealed that DeepVariant had lost more than half of its mutations (Figure 8, Supplementary Table S11).

**Figure 8:**
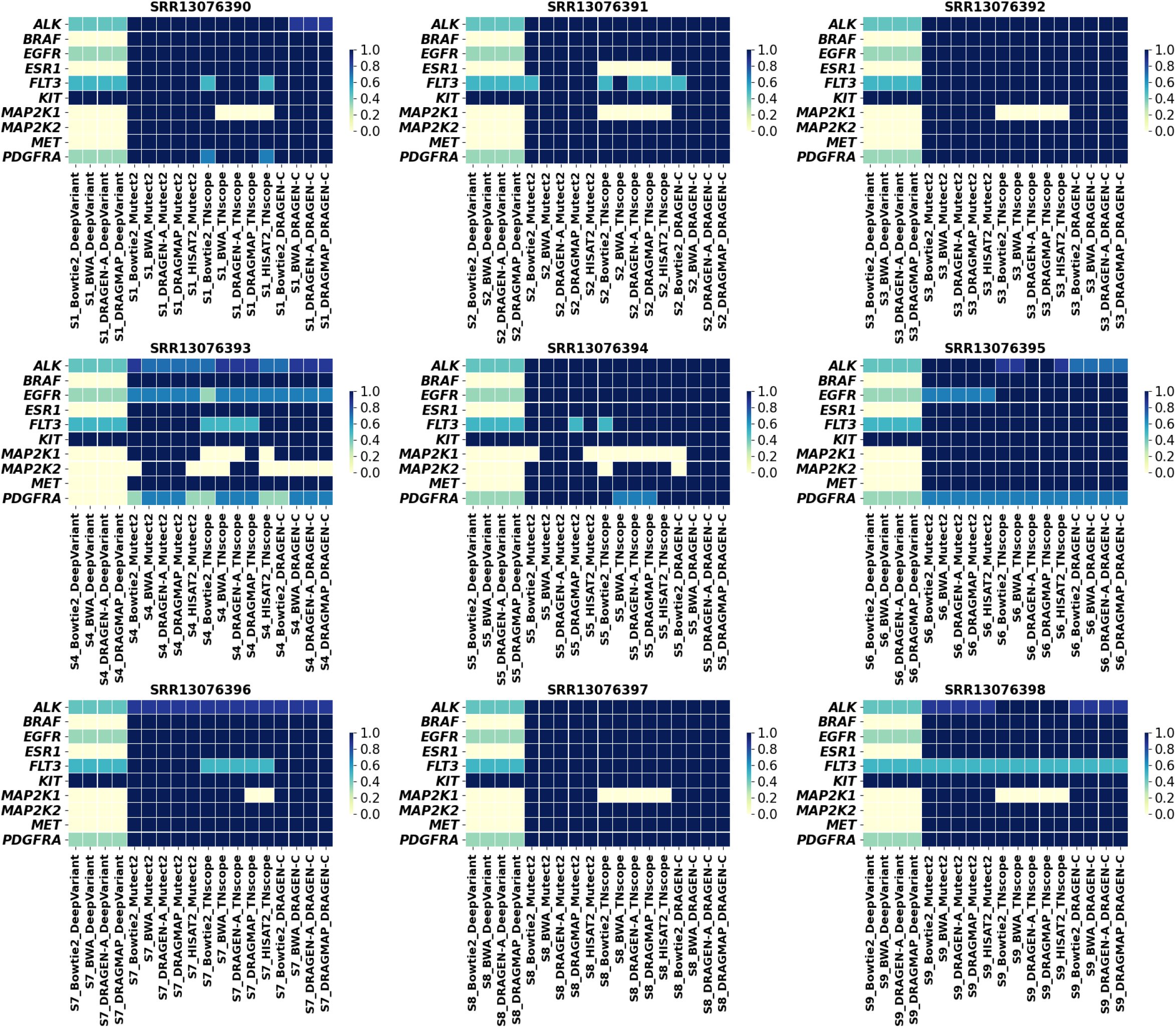
True positive discovery rate of the nine samples under different combinations in COSMIC-provided drug resistance variants.

Among all drug-associated mutations, mitogen-activated protein kinase 1, *MAP2K1* (c.G199A: p.D67N), is a key player in the mitogen-activated protein kinase (*MAPK*) signaling pathway that regulates cell growth, proliferation, and survival. Somatic mutations in *MAP2K1* have been identified in various cancers, including melanoma, colorectal cancer, and lung cancer. These mutations have garnered attention owing to their potential effect on treatment selection and drug responses. Studies have shown that patients harboring this mutation may exhibit resistance to specific targeted therapies, such as *BRAF* inhibitors, which are commonly used in melanoma treatment. This mutation has been shown to activate alternative signaling pathways that bypass the *MAPK* pathway, rendering tumor cells less responsive to *BRAF* - targeted treatments. All samples constructed using DeepVariant missed this variant and 36 of the 45 callsets constructed using TNscope failed to detect this variant. However, this mutation could be detected in sample 6 in all combinations except for the combination with DeepVariant.

Patients with FMS-like tyrosine kinase 3 (*FLT3* (c.G1879A:p.A627T)) mutations may exhibit increased resistance to conventional chemotherapy agents, such as cytarabine and anthracyclines. Therefore, alternative treatments are required. In addition to DeepVariant, TNscope tended to overlook this mutation. However, S3, S6, and S8 could detect *FLT3* (c.G1879A:p.A627T) by TNscope, suggesting that the mutation caller is not the only factor, but library preparation is also crucial. On the other hand, DRAGMAP Mutect2 combination, which has the best recall rate in SNV among open-source tools, also overlooked this mutation in sample S5. This example demonstrated that different tumor identification workflow can lead to distinct downstream clinical decisions. In summary, the study highlighted the technical importance of ensuring the reliability of tumor identification, encompassing both library preparation experiments and bioinformatic analysis workflows.

## 4. Discussion

Xiao et al. previously utilized breast cancer cell lines in comparison to matched normal cell lines to benchmark a multiple NGS-based tumor profiling workflow [5]. They addressed numerous experimental variables that influence the overall ability to detect somatic mutations, including biospecimen types, DNA input amounts, sequencing platforms across multiple centers, and library preparation methods. In this study, tumor-only data is concentrated on, which presents a more challenging and realistic scenario, as clinical settings often involve tumor tissue exclusively. Current commercial gene testings and retrospective research projects often use FFPE tumor samples only. Xiao et al demonstrated that WES exhibits greater variability in GIV scores, indicating that it is more sensitive to site-to-site library preparation variations than whole genome sequencing (WGS). The materials used in the study consist of a mixture of multiple well-characterized cell lines, enabling us to push the limits of detection for very low variant allele frequencies. While previous studies have targeted VAF as low as 5%, 33.4% of true positive calls in this study with VAF below 5%.

In this study, the performance of analysis workflows was evaluated using SEQC2 WES datasets for four popular short-read sequence aligners and five somatic mutation callers, resulting in 18 combinations for somatic mutation detection. The results showed that mutation callers had a notably higher effect on the overall sensitivity than read aligners. For the overall aligner-caller combinations, the Illumina DRAGEN built-in DRAGEN-A DRAGEN-C combination had the highest F1 scores for both SNVs and INDELs, followed by BWA Mutect2 and HISAT2 Mutect2, which were the best open-source combinations for SNVs and INDELs, respectively. DRAGEN showed the best performance, with a mean F1-score of 0.9659 for SNV detection, whereas the combination of BWA and Mutect2 showed the second-highest mean F1-score of 0.9485. A more comprehensive targeted region with a lower duplication rate is recommended for library preparation. Even though library preparation is crucial, lab protocols can significantly impact performance beyond the library preparation reagents itself. For imp roved recall in SNV, DRAGEN-A DRAGEN-C and DRAGMAP Mutect2 are suggested as open-source option. For recall in INDEL, DRAGMAP DRAGENC, with DRAGMAP Mutect2 are recommend as the open-source alternative. To enhance precision in SNV, BWA TNscope and BWA Mutect2 are advised for the open-source option. For precision in INDEL, HISAT2 TNscope and HISAT2 Mutect2 are suggested as open-source alternatives. The somatic VAF is a critical factor in somatic mutation analyses and is different from germline variant detection. The VAF may influence the variant detection capability in different combinations. Some general variant detection tools, such as DeepVariant, perform inadequately. The analysis results also suggest that sequencing library preparation kits influence mutation detection capabilities, as Agilent library preparation kits have overall higher F1 scores. The results of the shared variants in different combinations showed that even if some combinations demonstrated superiority in the F1-scores, there were still several variants that were not detected. These FN mutations may be important because of their implications for personalized therapeutic strategies for patients. The data demonstrated that Sentieon TNscope often underestimated the TMB and missed drug-resistance mutations, such as *FLT3* (c.G1879A:p.A627T) and *MAP2K1* (c.G199A:p.D67N). While this study only relied on single benchmarking dataset, which is a limitation, it provides a valuable guide for cancer genomic researchers in tumor mutation identification. Even though several other excellent short-reads aligners and somatic mutation callers are available, the selected tools in this study are all state-of-the-art. This study also accomplished an in-depth performance comparison among diverse tool combinations. Current clinical cancer panels often focus on a group of cancer-associated genes and the sequencing depth can reach at least 1000 folds. However, WES might be used extensively with increasing number of genes identified to be associated with cancer.

The results of this study provide supportive evidence that the ensemble-based approach using multiple variant callers or combinations can improve the results of variant detection using one or few variant callers or combinations. Further benchmark analyses are necessary during clinical genetic testing.

## 5. Author contributions

PeiMiao Chien analyzed data, visualized the results, and contributed in writing the first draft manuscript. Chinyi Cheng initiated the data analyses, interpreted results, and contributed in writing the first draft manuscript. Larry Lin analyzed the sequence data and curated the results. Tzu-Hang Yuan and Yu-Bin Wang analyzed the sequence data. Pei-Lung Chen conceptualized the project and curated the data. Chien-Yu Chen and Jia-Hsin Huang conceptualized and supervised the project. Jacob Shujui Hsu conceptualized and supervised the project, acquired funding, and contributed to writing the manuscript. All authors read and approved the final manuscript.

## 6. Acknowledgments

Appreciation is extended to all technicians and investigators who contributed to this study. Gratitude is also extended to the National Center for High-performance Computing (NCHC) for providing computational and storage resources. This study was supported by the research grants (NSTC 111-2320-B-002-091-MY3, 112-2320-B-002-045) and industry-academia cooperative research program grants (NSTC 110-2622-B-002-012, 111-2622-B-002-012, 112-2622-B-002-012) from the National Science and Technology Council in Taiwan. This project is also supported by research grants from National Taiwan University (113L892901).

