## Supplementary material for "Evaluating discordant somatic calls across mutation discovery approaches to minimize false negative drug-resistant findings": supply (from another version of this preprint: version 2, 2024-11-19)

**Supplementary Figures**

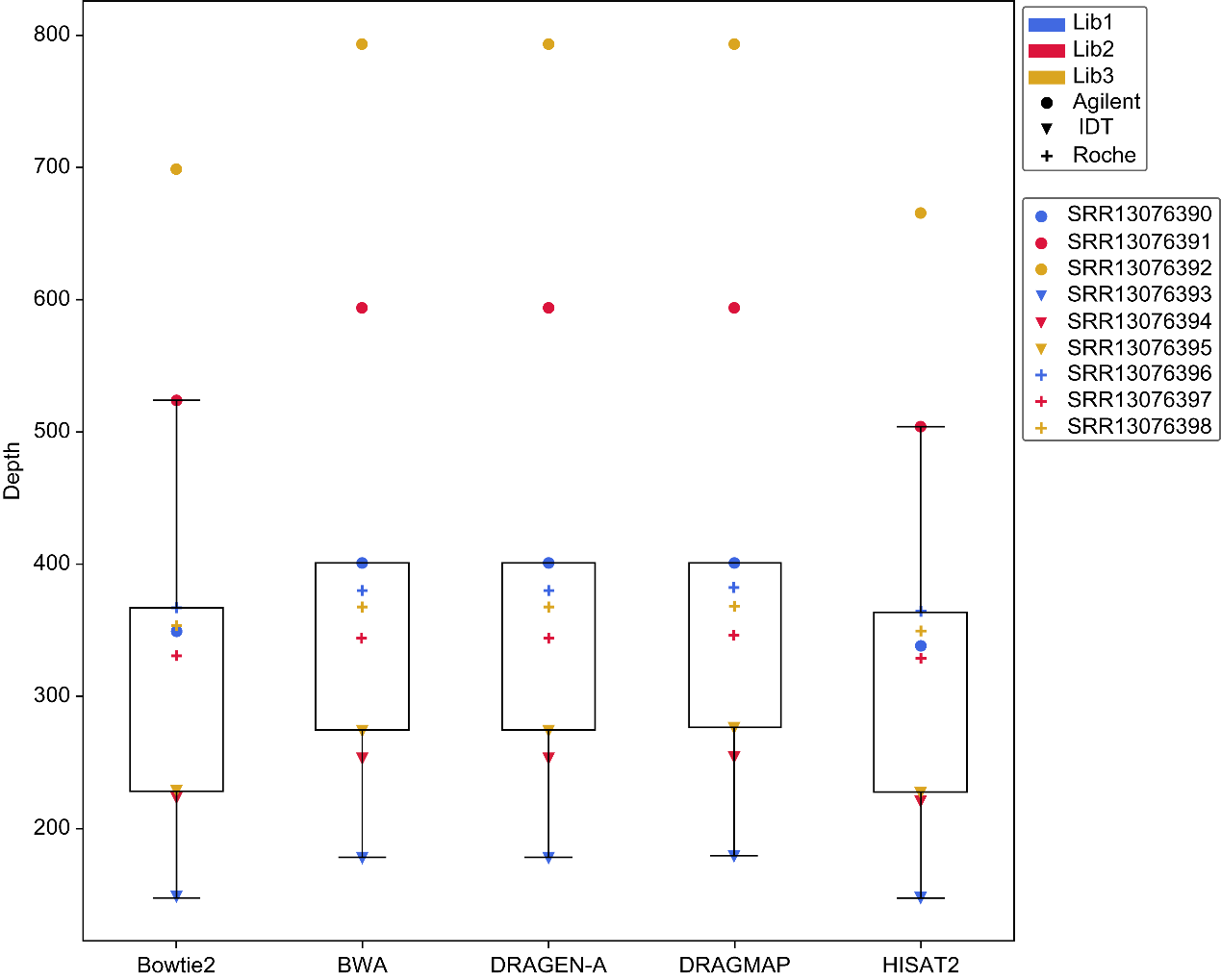

### Supplementary Figure S1. Comparison of the overall deduplicated mapping depths of nine WES samples among three laboratories and three library preparation kits (Agilent/IDT/Roche) under different aligners.

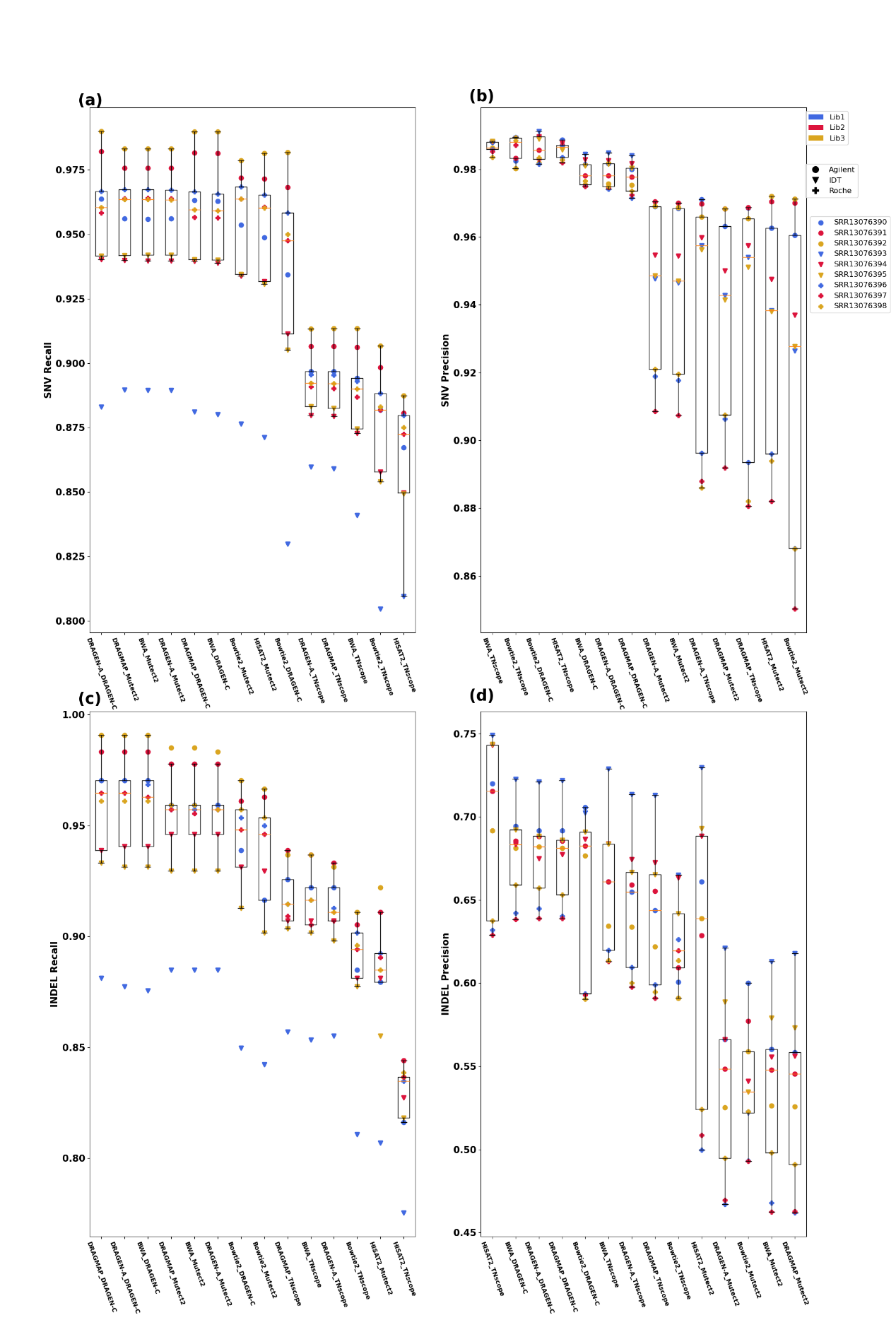

### Supplementary Figure S2. (a) and (b) showed comparison of the recall and precision of somatic mutation calls among workflows with SEQC2 WES datasets as a truth set for SNV. (c) and (d) showed comparison of the recall and precision of somatic mutation calls among workflows with SEQC2 WES datasets as a truth set for INDEL.

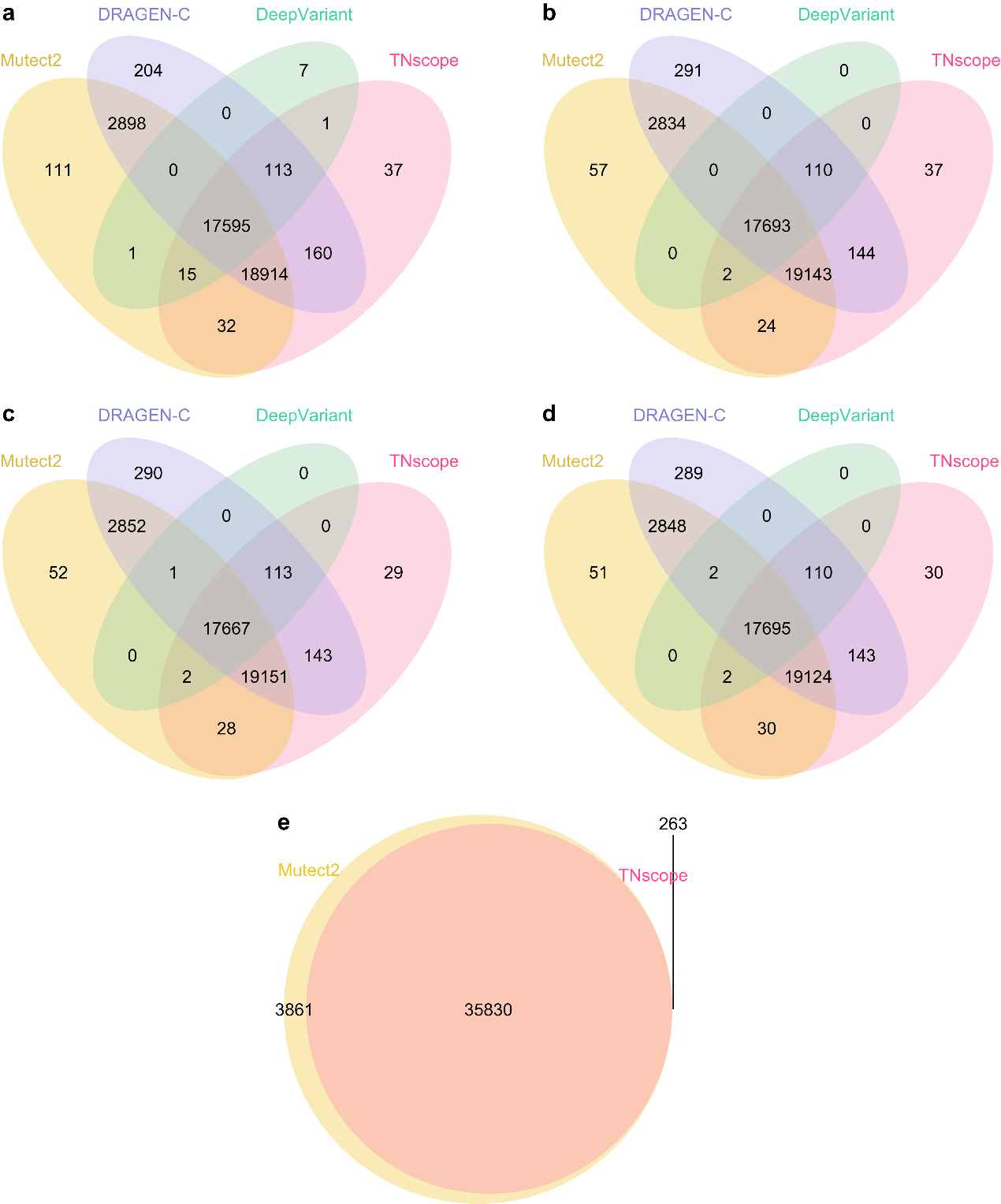

### Supplementary Figure S3. The number of SNV calls from the same aligner among variant callers. (A) Bowtie2, (B) BWA, (C) DRAGEN-A, (D) DRAGMAP, and (E) HISAT2.

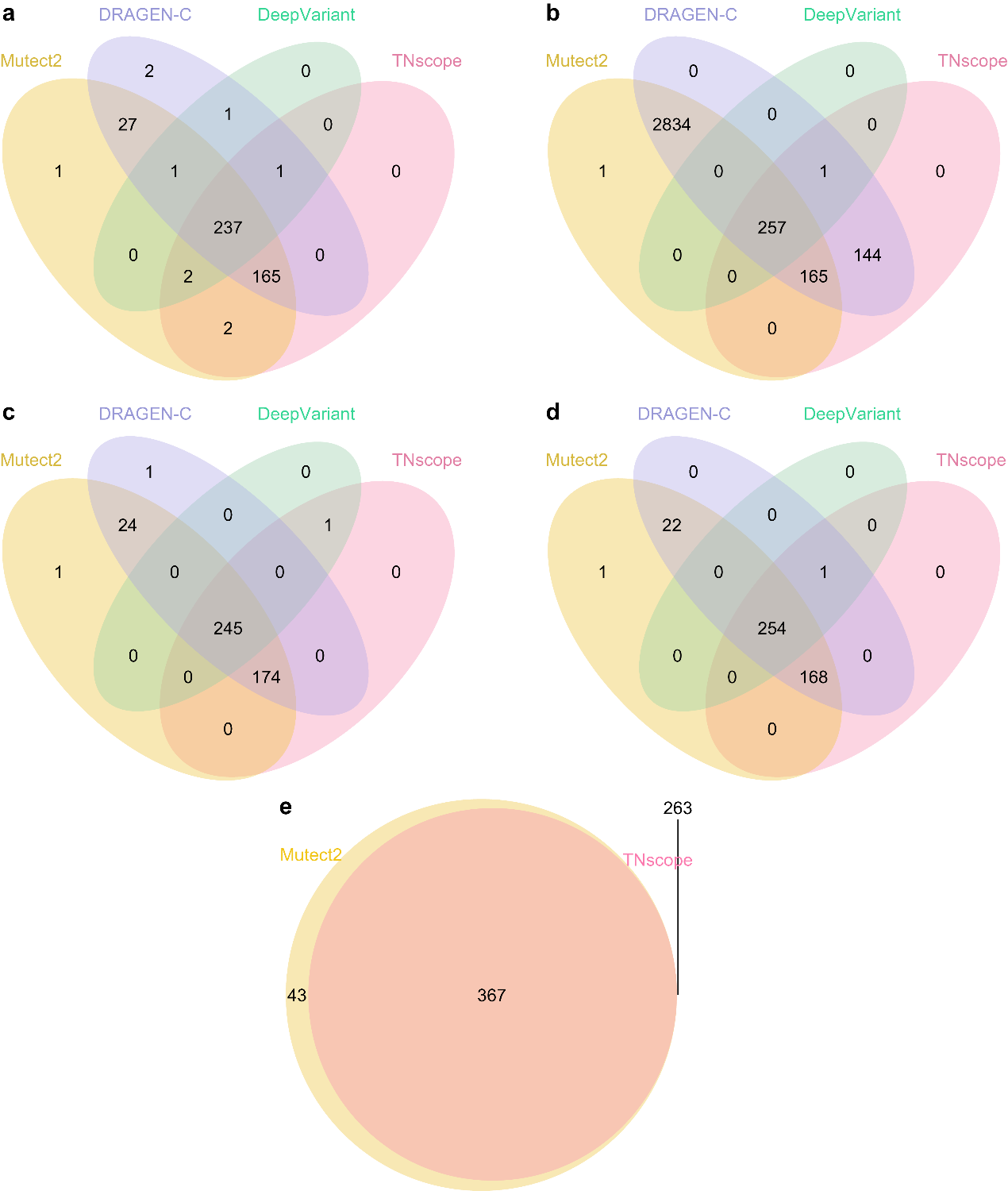

### Supplementary Figure S4. The number of INDEL calls from the same aligner among variant callers. (A) Bowtie, (B) BWA, (C) DRAGEN-A, (D) DRAGMAP, and (E) HISAT2.

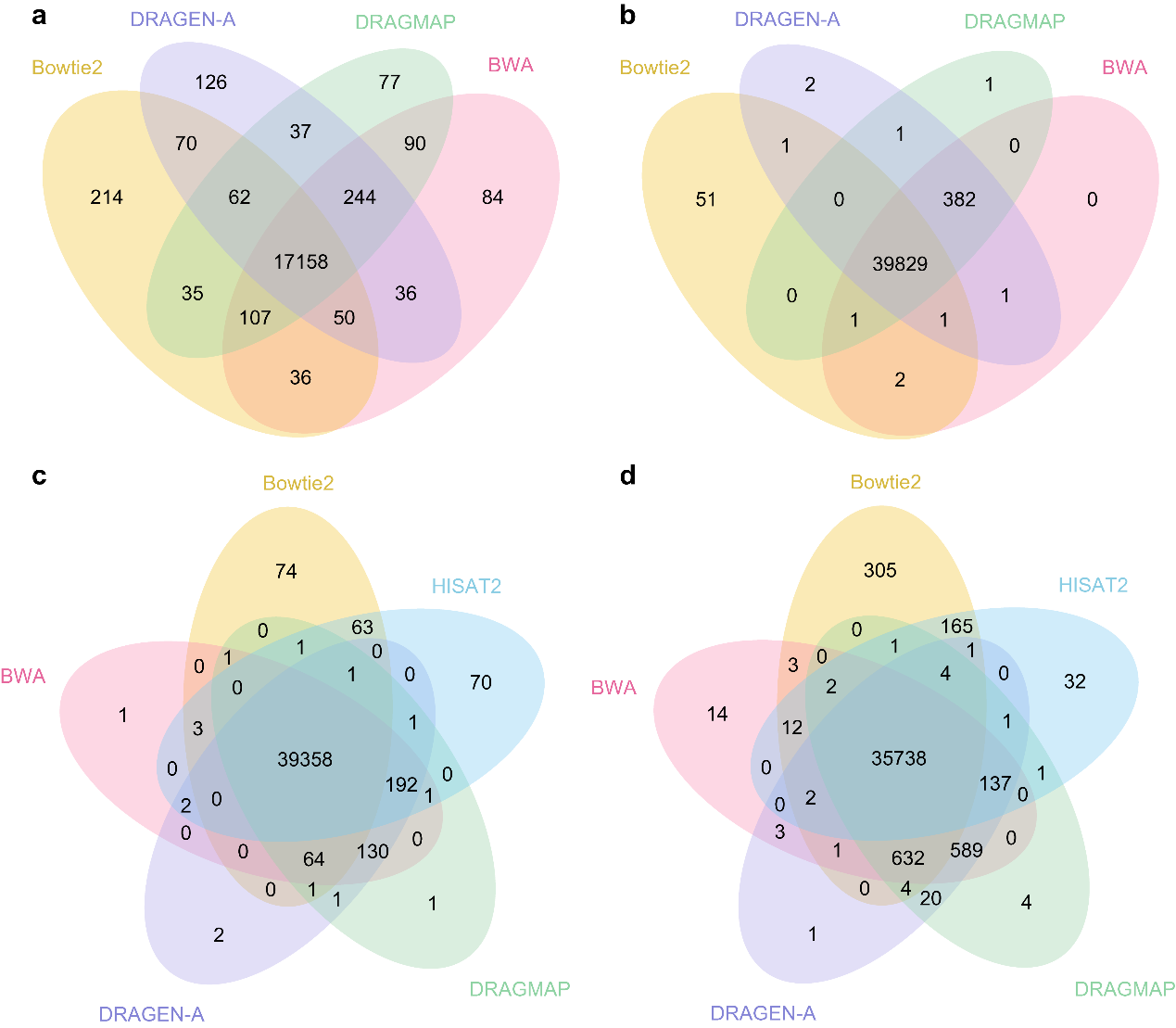

### Supplementary Figure S5. SNV calling results of the different aligner and same caller: (A) DeepVariant, (B) DRAGEN-C, (C) Mutect2, and (D) TNscope.

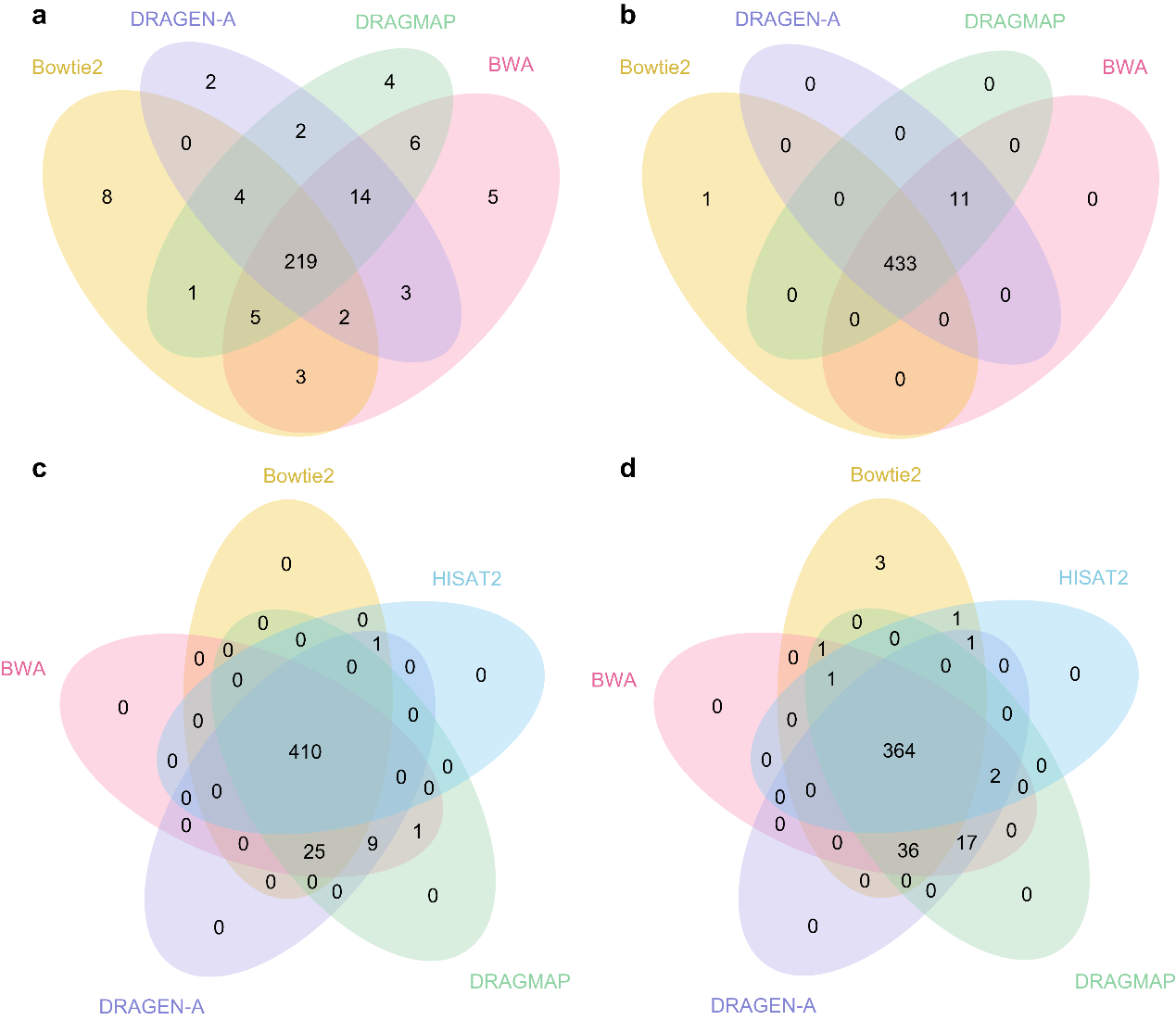

### Supplementary Figure S6. INDEL calling results of the different aligner and same caller: (A) DeepVariant, (B) DRAGEN-C, (C) Mutect2, and (D) TNscope.

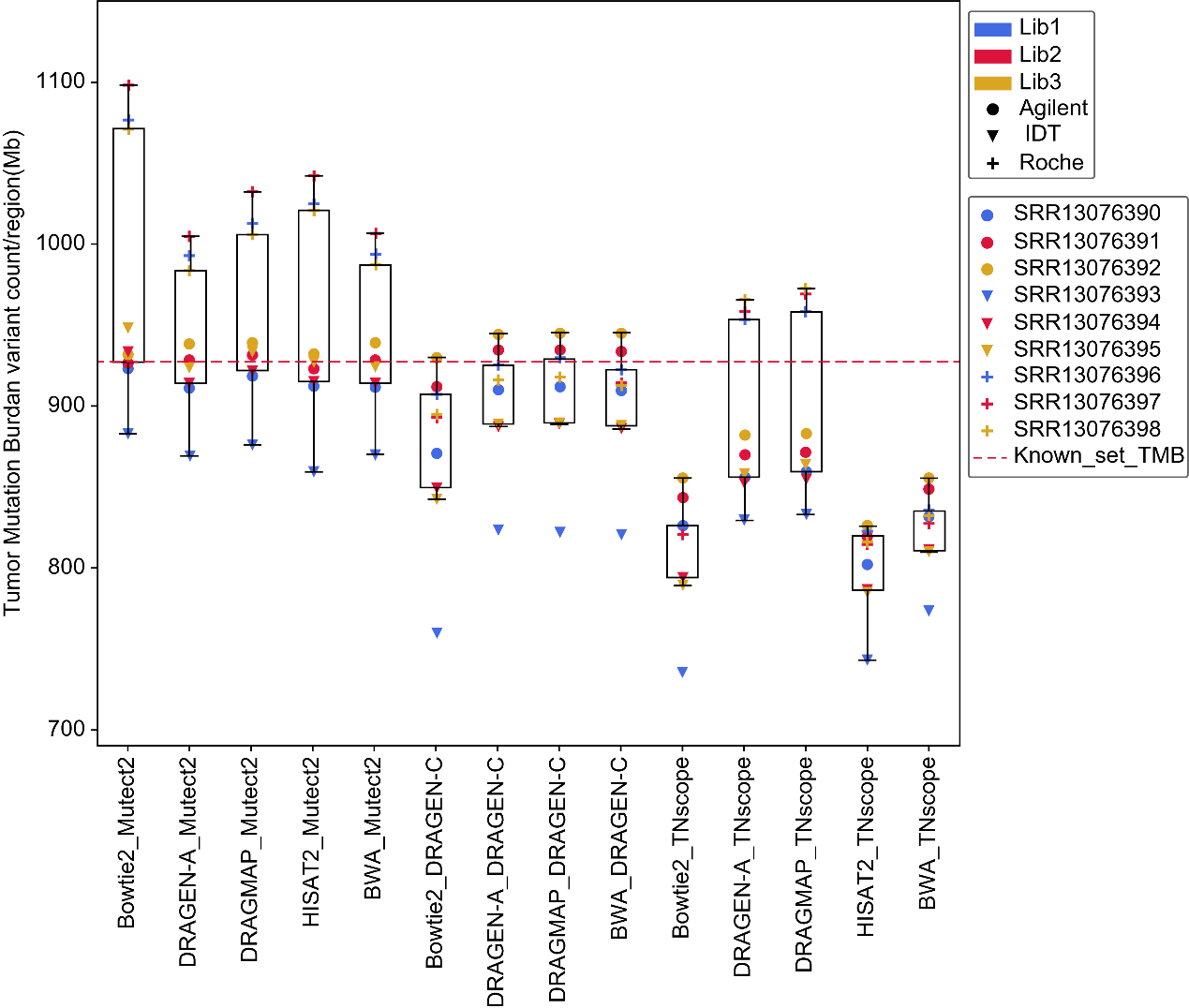

### Supplementary Figure S7. Comparison of the tumor mutation burden (TMB) results among combinations for all nine SEQC2 samples. The red line indicates the truth set TMB.

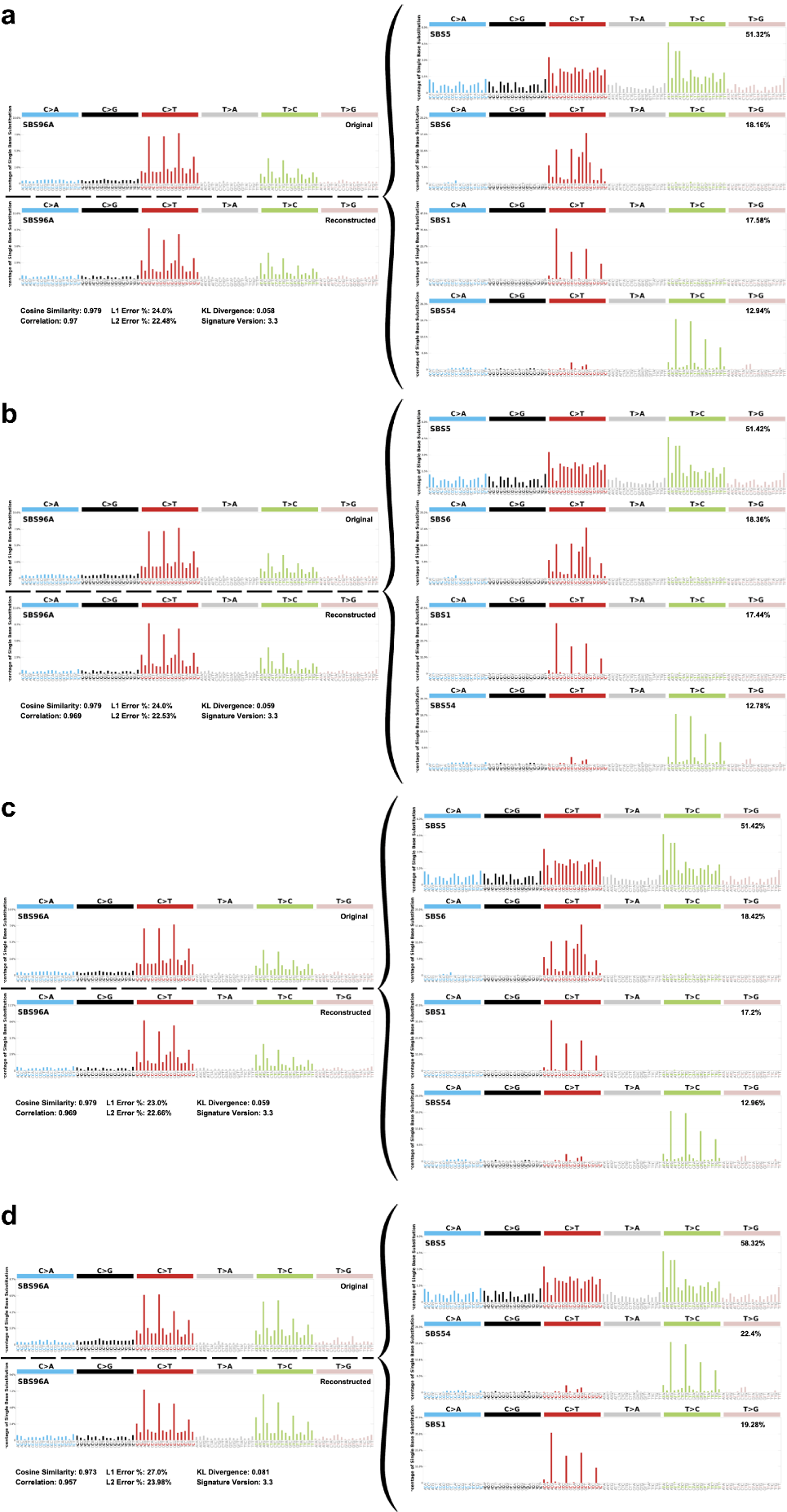

**Supplementary Figure S8.** The single-base mutational signatures (SBS) of sample S3 (SRR13076392) from four tool combinations: (A) Known sets, (B) DRAGEN-A_DRAGEN-C, (C) BWA_Mutect2, and (D) Bowtie2_DeepVariant.

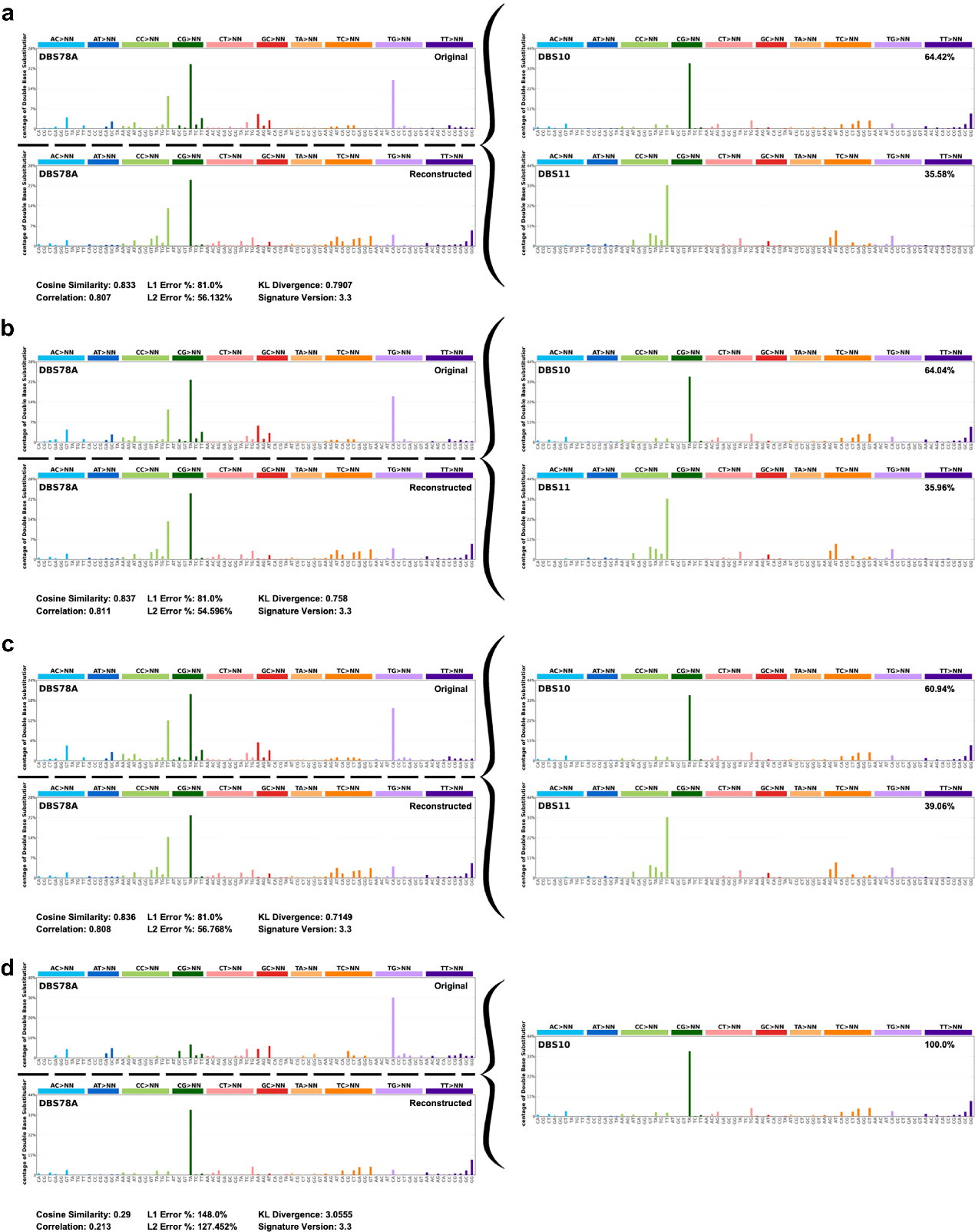

**Supplementary Figure S9.** The double-base mutational signatures (DBS) of sample S3 (SRR13076392) from four tool combinations: (A) Known sets, (B) DRAGEN-A_DRAGEN-C, (C) BWA_Mutect2, and (D) Bowtie2_DeepVariant.

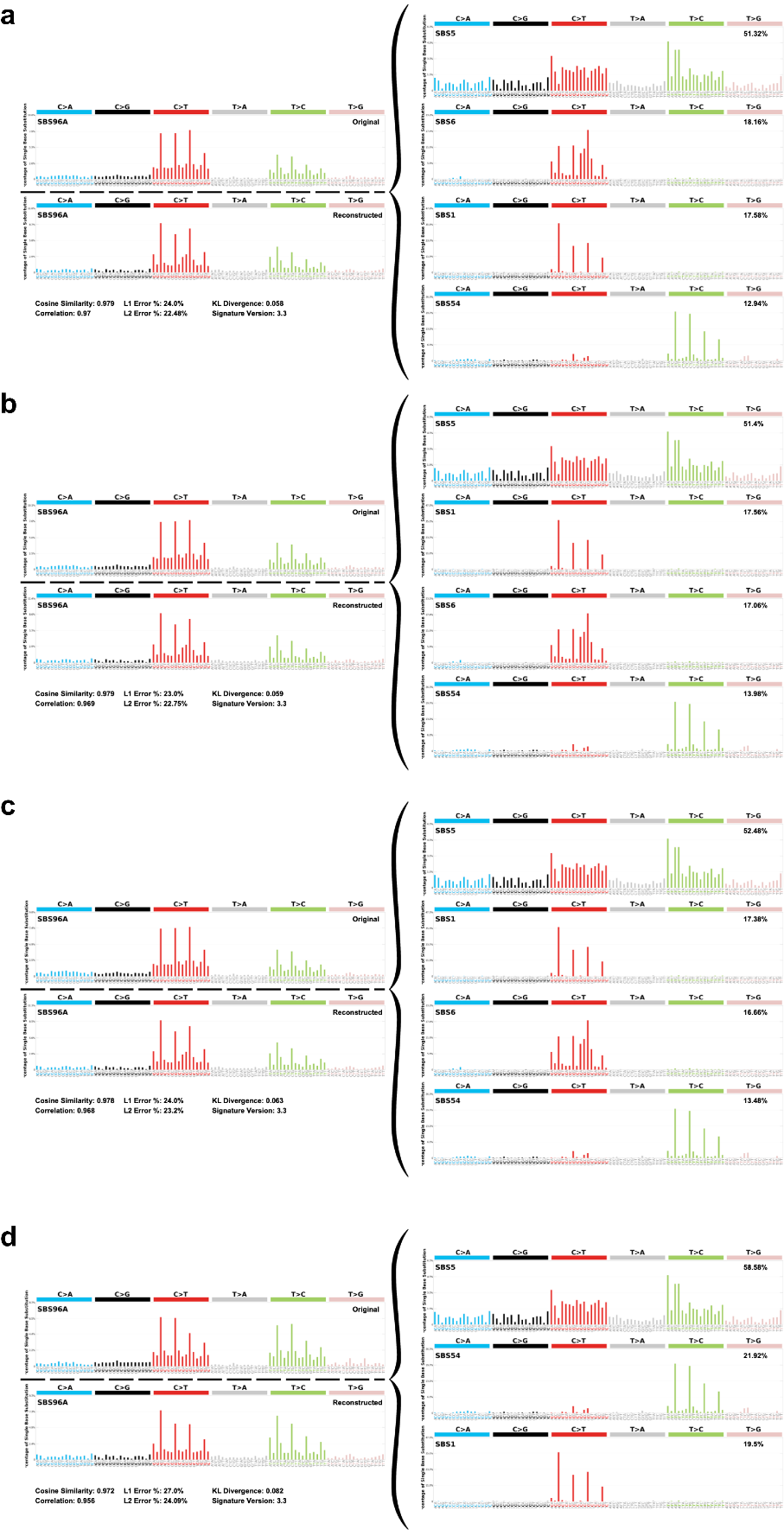

**Supplementary Figure S10.** The single-base mutational signatures (SBS) of sample S4 (SRR13076393) from four tool combinations: (A) Known sets, (B) DRAGEN-A_DRAGEN-C, (C) BWA_Mutect2, and (D) Bowtie2_DeepVariant.

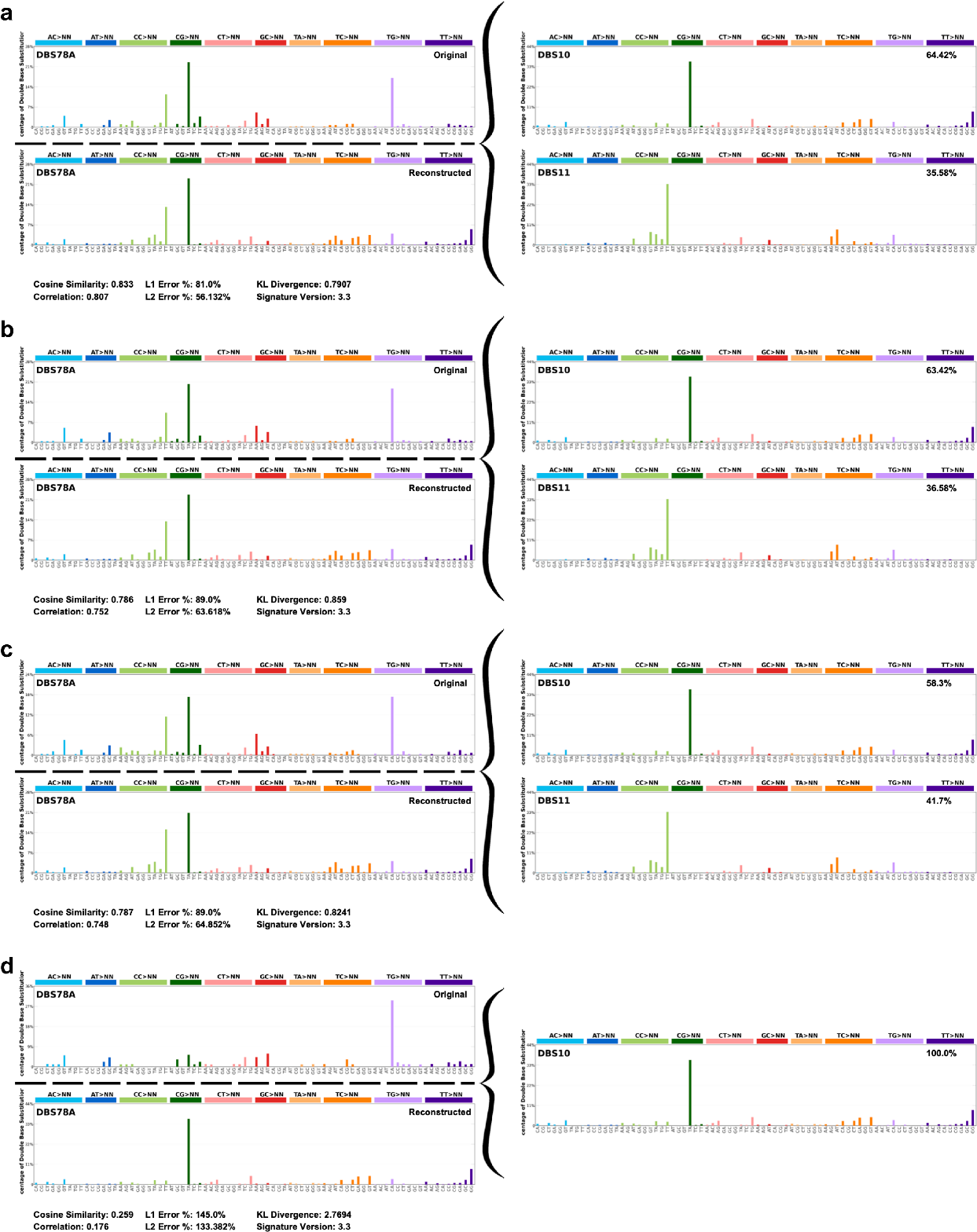

**Supplementary Figure S11.** The double-base mutational signatures (DBS) of sample S4 (SRR13076393) from four tool combinations: (A) Known sets, (B) DRAGEN-A_DRAGEN-C, (C) BWA_Mutect2, and (D) Bowtie2_DeepVariant.

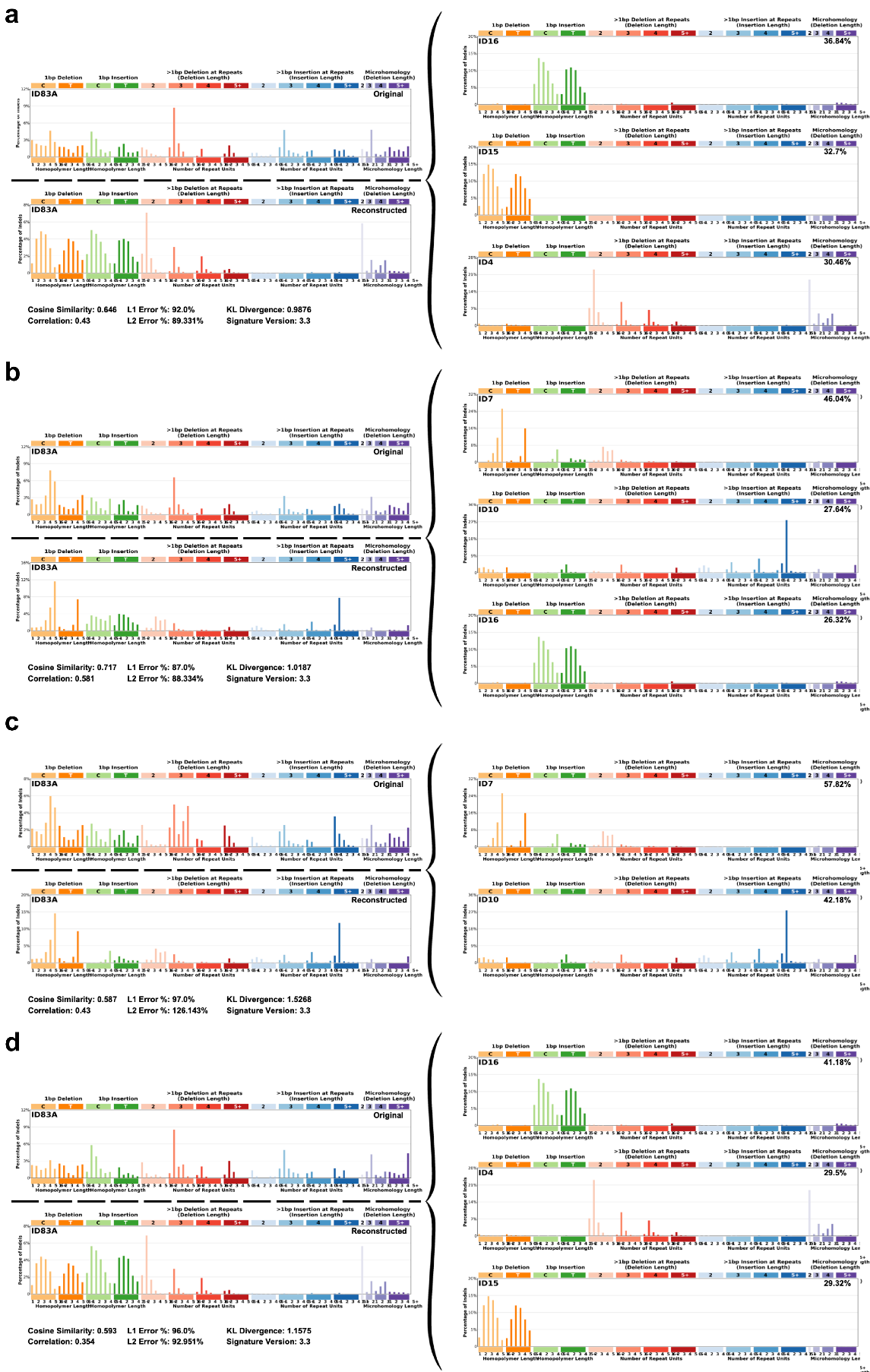

**Supplementary Figure S12.** The INDEL mutational signatures (ID) of sample S3 (SRR13076392) from four tool combinations: (A) Known sets, (B) DRAGEN-A_DRAGEN-C, (C) BWA_Mutect2, and (D) Bowtie2_DeepVariant.

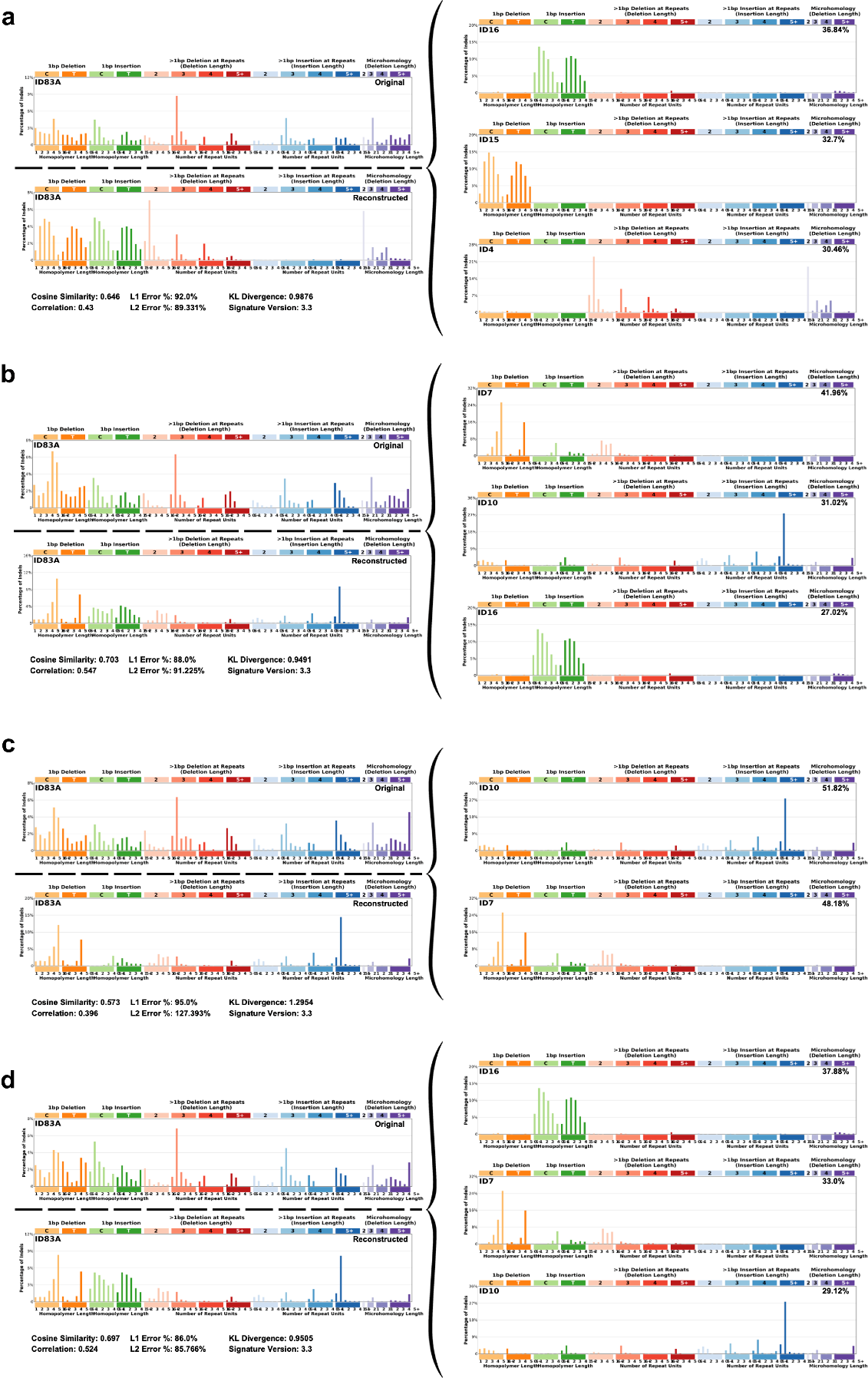

**Supplementary Figure S13.** The INDEL mutational signatures (ID) of sample S4 (SRR13076393) from four tool combinations: (A) Known sets, (B) DRAGEN-A_DRAGEN-C, (C) BWA_Mutect2, and (D) Bowtie2_DeepVariant.

The COSMIC list of mutational signatures was also inspected, including single-base signatures (SBS), double-base signatures (DBS), and INDEL signatures (ID). Table 2 shows a composition of the mutational signatures based on data from the best combination, DRAGEN, the best open-source combination, BWA_Mutect2, and the worst combination, Bowtie2_DeepVariant. All results (Supplementary Table 10) across different library preparation kits and combinations for SBS analyses shared similar patterns with the truth set, except for DeepVariant. The Bowtie2_DeepVariant combination failed to detect SBS6 in samples S3 (SRR13076392) and S4 (SRR13076393). SBS6 is linked to microsatellite instability (MSI) because of defective DNA mismatch repair. In the DBS analyses, Bowtie2_DeepVariant lost DBS11, which has been suggested to be associated with APOBEC mutagenesis. In ID analyses, even DRAGEN for sample 3 could recover just one mutational signature (ID6), indicating the systemic technical errors and irreproducibility of INDEL results (Supplementary Fig. 8-13).
